# Seasonal climatic impacts on orchid productivity in an urban ecosystem

**DOI:** 10.64898/2026.06.29.735162

**Authors:** Mark C. Brundrett

## Abstract

**Context:** The global diversity hotspot in Southwest Australia has >480 orchids facing increasing threats from climate extremes, fire and habitat decline.

**Aims:** To develop effective and consistent tools for measuring climate impacts on productivity in a diverse urban orchid community.

**Methods:** Annual variations in flower and seed production for 17 orchids were determined using thousands of records over a decade with extreme climate variability.

**Key results:** Rainfall deficits and temperatures in autumn, winter and spring increased substantially over 125 years. Seasonal climate anomalies reduced flowering and seed production for orchids, but this varied between species and seasons. These effects were summarised by climate response (CRI) and sensitivity (CSI) indexes. Early or late flowering species were most vulnerable to seasonal drought, but warm dry conditions promoted visually deceptive pollination. CRIs were strongly correlated with orchid pollination syndromes and flowering times. Effects on mycorrhizal fungi and pollinators were also observed. Extrapolating climate trends to 2100 predicted further impacts on orchid productivity (5-40%).

**Conclusions:** Orchid climate responses were substantial, complex and deeply integrated with key traits such as pollination, phenology and fire responses.

**Implications:** Research in an urban climate observatory produced a climate analysis framework potentially relevant to all orchids and other biota.

## Introduction

The Southwest Australian Floristic Region (SWAFR) is a major global diversity hotspot for plant species richness and endemism (Myers et al. 2000, Hopper & Gioia 2004). Plant functional and taxonomic diversity in this region is linked to long-term landscape stability, complexity and extreme infertility (Groom & Lamont 2015, Brundrett 2021). Plant functional diversity includes exceptionally functional trait complexity, with 90% having multiple fire survival adaptations (Brundrett 2021) and over 275 pollination trait transitions (Brundrett et al 2024). The SWAFR also has an exceptional diverse terrestrial orchid flora (487 taxa, florabase.dbca.wa.gov.au, accessed 23-6-2026). These orchids face increasing impacts from a drying and warming climate, fire, weeds, grazing, disturbance, etc. (Duncan et al. 2005, Brundrett 2014, Martín-Forés et al. 2022, Doherty et al. 2024). However, orchid biology also provides advantages such as wind dispersed seeds and drought avoidance by vegetative dormancy (Brundrett 2007). For West Australian (WA) orchids, biological factors that regulate population sustainability include very specific pollinators attracted by visual (44%) or sexual (38%) deception (Brundrett et al. 2024) and extreme nutritional dependencies on mycorrhizal fungi, due to sparse or absent roots (Ramsay et al. 1986; Brundrett 2014).

Substantial rainfall declines and temperature increases have occurred in southwest Western Australia in the past century (Scanlon et al. 2020, Verhoeven et al. 2022, www.bom.gov.au), but need to be considered in the context of climate history reconstructions which show decadal cycles of drought and wet periods over centuries (O’Donnell et al. 2018). Despite these complexities, it has been established that recent climate trends have had substantial impacts on Australia vegetation, and have strong fire, grazing and disturbance interactions (Hoffman et al. 2019, Williams et al. 2025). Vegetation decline in both urban woodlands and large forest areas also result from extreme temperatures and groundwater drawdown linked to rainfall deficits and over-extraction (Groom et al. 2000, Froend & Sommer 2010, McFarlane et al 2025, Wang et al. 2025).

Herbarium studies show that climatic change has altered orchid flowering times and reduced rates of pollen removal or deposition, especially in the past 50 years (Gallagher et al. 2009, Robbirt et al. 2011, Molnár et al. 2012, Song et al. 2025). However, data on direct climatic impacts on orchid productivity and reproduction are limited to a few case studies. These include, reduced flowering due to high temperatures for *Prasophyllum tadgellianum,* but not *Prasophyllum suttonii* (Gallagher et al. 2009), reduced seed production with low rainfall for *Diuris maculata* (Brown and York 2016), rainfall effects on flowering of *Pterostylis revoluta* (Jasinge et al. 2018), and benefits from increased rainfall and longer growing seasons for a rare *Caladenia* species (Janissen 2021). Observations in WA link increased rainfall and pollination for *Leporella fimbria*, a sexually deceptive autumn-flowering orchid, while dry warm weather increased capsule production for visually deceptive *Thelymitra* species (Brundrett 2019), and emergence and flowering were strongly dependant on rainfall for *Caladenia graniticola,* a rare orchid from relatively dry habitats (Brundrett 2016).

Banksia woodlands are threatened plant communities centred around the Perth Metropolitan Region on the Swan Coastal Plain in Western Australia (Commonwealth of Australia 2016). This bioregion is impacted by extensive land clearing and habitat fragmentation, as well as increasing impacts from fire, weeds and climate change (Ramalho et al. 2014; Ritchie et al. 2021, Fowler et al. 2023, Fay et al. 2025). It also has a very diverse flora including over 150 species of indigenous terrestrial orchids (Brundrett & George in prep.). This study focussed on an isolated urban remnant of banksia woodland which is a local orchid diversity hotspot. Specific objectives were to: (i) quantify long-term trends and recent seasonal fluctuations in climatic conditions, (ii) develop climate tolerance indexes based on orchid productivity data, (iii) compare seasonal variations in climate sensitivity between species and their correlations with other traits, (iv) separate direct effects on orchids from indirect effects on insects or fungi, (v) select appropriate indicator species, and (vi) develop a climate monitoring toolkit.

## Methods

### 1. Study site

This study occurred in Warwick Bushland, an urban reserve of 63 hectares, located in the City of Joondalup 13 km north of the centre of Perth in Western Australia. This Bush Forever Site is protected due its size, biodiversity and vegetation condition (Government of Western Australia 2000). Comprehensive information on biodiversity, history and fire is available (friendsofwarwickbushland.com, accessed 13-6-2026). There are large areas of very good or excellent condition vegetation in this isolated area (∼5 km from similar habitats). The 217 native flora species include 30 orchids, of which 12 are regionally significant. Vegetation is dominated by *Banksia* species with a scattered overstory of eucalypts. Dominant trees are *Banksia attenuata*, *B. menziesii* and *Eucalyptus marginata*, with scattered *Allocasuarina fraseriana* and *Eucalyptus gomphocephala*. Vegetation is in the Karrakatta Central and South Complex of the Spearwood Dune system. Soils are highly infertile grey sands around 40,000 years old (Government of Western Australia 2000).

### 2. Study species

This study focused on orchids included in ongoing detailed pollination and fire research (Brundrett 2019, 2025) using consistent observations of plants in Warwick Bushland (Table S1, Table 1). Two common highly clonal species with intermittent fire-induced flowering were omitted here due to infrequent flowering (*Pyrorchis nigricans* and *Leptoceras menziesii).* The remaining 17 species had 7 to 10 years of productivity data totalling over 30,000 observations (Table S1). Pollination strategies, typical rates of flower production and pollination, fire responses and other key traits of these species are summarised in Table 1.

**Table 1.** Orchids included in study with their size, ecology, phenology and productivity (2016-2025)

|  |  | Pollination |  |  | Width, height, length |  |  |  | Phenology dates |  |  | Average values |  | Capsule | Study | Total records for decade |  |  |  |
| --- | --- | --- | --- | --- | --- | --- | --- | --- | --- | --- | --- | --- | --- | --- | --- | --- | --- | --- | --- |
| Orchid | Clade | Label | syndrome | Flowering | Flower | Spike | Leaf (no.) | Records | Leaf | Flowers | Seed | Flowers | Pollination | volume | period | Plants | Spikes | Flowers | Capsules |
| *Caladenia arenicola* | Caladeniinae* | Ca | SD wasp | spring | 58 | 450 | 200 | 64 | 185 | 259 | 281.5 | 1.4 | 16% | 360 | 10 | 588 | 529 | 729 | 117 |
| *Caladenia discoidea* | Caladeniinae* | Cd | SD wasp | spring | 25 | 220 | 130 | 11 | 196 | 255 | 263.5 | 1.5 | 35% | 350 | 10 | 125 | 122 | 178 | 55 |
| *Caladenia flava* | Caladeniinae* | Cf | VD | spring | 33 | 175 | 90 | 37 | 182 | 250 | 262 | 2.0 | 24% | 280 | 9 | 1924 | 719 | 1407 | 219 |
| *Caladenia latifolia* | Caladeniinae* | Cl | VD | spring | 30 | 325 | 175 | 27 | 187 | 252 | 262.5 | 2.4 | 34% | 320 | 9 | 1174 | 360 | 859 | 405 |
| *Disa bracteata* | Diseae (Africa) | Db | Selfing | late spring | 10 | 300 | 90 (x9) | 30 | 205 | 314 | 316.5 | 26.0 | 55% | 72 | 9 | 778 | 776 | 20190 | 12293 |
| *Diuris magnifica* | Diuridinae* | Dm | VD | spring | 35 | 425 | 225 | 48 | 177 | 247.5 | 262 | 3.7 | 5% | 320 | 10 | 5456 | 4608 | 17085 | 878 |
| *Elythranthera brunonis* | Caladeniinae* | Eb | VD | spring | 28 | 225 | 85 | 33 | 183 | 259.5 | 271.5 | 1.8 | 18% | 180 | 10 | 428 | 331 | 584 | 85 |
| *Leporella fimbria* | Megastylidinae* | Lf | SD ant | autumn | 20 | 175 | 25 | 76 | 142 | 137 | 152.5 | 1.7 | 32% | 45 | 10 | 10984 | 1719 | 2962 | 980 |
| *Microtis media* | Prasophyllinae* | Mm | Selfing | late spring | 4 | 450 | 450 | 27 | 205 | 308 | 316.5 | 34.4 | 88% | 14 | 9 | 1029 | 1029 | 35363 | 33501 |
| *Pheladenia deformis* | Caladeniinae* | Pd | VD | Winter | 20 | 100 | 65 | 39 | 155 | 187.5 | 219 | 1.0 | 10% | 190 | 8 | 1247 | 829 | 830 | 94 |
| *Pterostylis ectypha* | Cranichideae | Pe | SD gnat | Winter | 9 | 75 | 25 (x6) | 34 | 193 | 224 | 237.5 | 1.0 | 48% | 80 | 7 | 1223 | 909 | 909 | 433 |
| *Pterostylis orbiculata* | Cranichideae | Po | SD gnat | Winter | 18 | 350 | 30 (x14) | 47 | 133 | 170.5 | 192.5 | 4.0 | 40% | 300 | 8 | 2647 | 1789 | 7125 | 2976 |
| *Thelymitra benthamiana* | Thelymitrininae* | Tb | Selfing | spring | 33 | 300 | 150 | 27 | 191 | 290.5 | 310 | 2.4 | 88% | 350 | 9 | 1108 | 488 | 1175 | 1045 |
| *Thelymitra fuscolutea* | Thelymitrininae* | Tf | VD | late spring | 33 | 300 | 100 | 62 | 210 | 327.5 | 336.5 | 6.2 | 7% | 160 | 10 | 2021 | 868 | 5408 | 312 |
| *Thelymitra graminea* | Thelymitrininae* | Tg | VD | spring | 25 | 200 | 105 | 14 | 191 | 293 | 305 | 4.6 | 16% | 140 | 9 | 389 | 384 | 1766 | 240 |
| *Thelymitra macrophylla* | Thelymitrininae* | Tm | VD | spring | 40 | 500 | 200 | 47 | 142 | 272.5 | 289.5 | 13.9 | 4% | 215 | 10 | 9861 | 5768 | 80288 | 2901 |
| *Thelymitra vulgaris* | Thelymitrininae* | Tv | Selfing | spring | 18 | 200 | 175 | 15 | 180 | 277 | 272.5 | 2.7 | 93% | 133 | 9 | 154 | 127 | 345 | 319 |
| Units | SD = sexual deception, VD = visual deception |  |  |  | mm | mm | mm (ave.) | no. | middle growth period (d) |  |  | per spike | per flower | mm<sup>3</sup> | years | no. | no. | no. | no. |
| **Total or Average** | *Diuridaea |  |  |  | **26** | **281** | **155** | **638** | **180** | **254** | **268** | **6.5** | **36%** | **206** | **9.2** | **41,136** | **21,355** | **177,203** | **56,853** |

### 3. Climatic data

Long-term rainfall and temperature data from the Bureau of Meteorology (BOM, www.bom.gov.au) were amalgamated from several weather stations close to the study site to make a continuous 149-year record for rainfall or 118 years for temperature to the end of 2025 (Fig. 1). These records were also compared with each other (Fig. 2). Rainfall data from a private rain gauge < 1 km from the site and the BOM Wanneroo weather station 11 km away were in close overall agreement, but had gaps so were consolidated into a continuous record (Table S2, Fig. 1). Rainfall totals in autumn, winter and spring for the study period (2016-2025), were summarised and converted to rainfall anomalies using long-term averages (Fig. 3). Dates for the start and end of growing seasons were determined from daily rainfall data (Pook et al. 2009) and used to calculate growing season length, initiation and end dates during the study.

**Figure 1.**
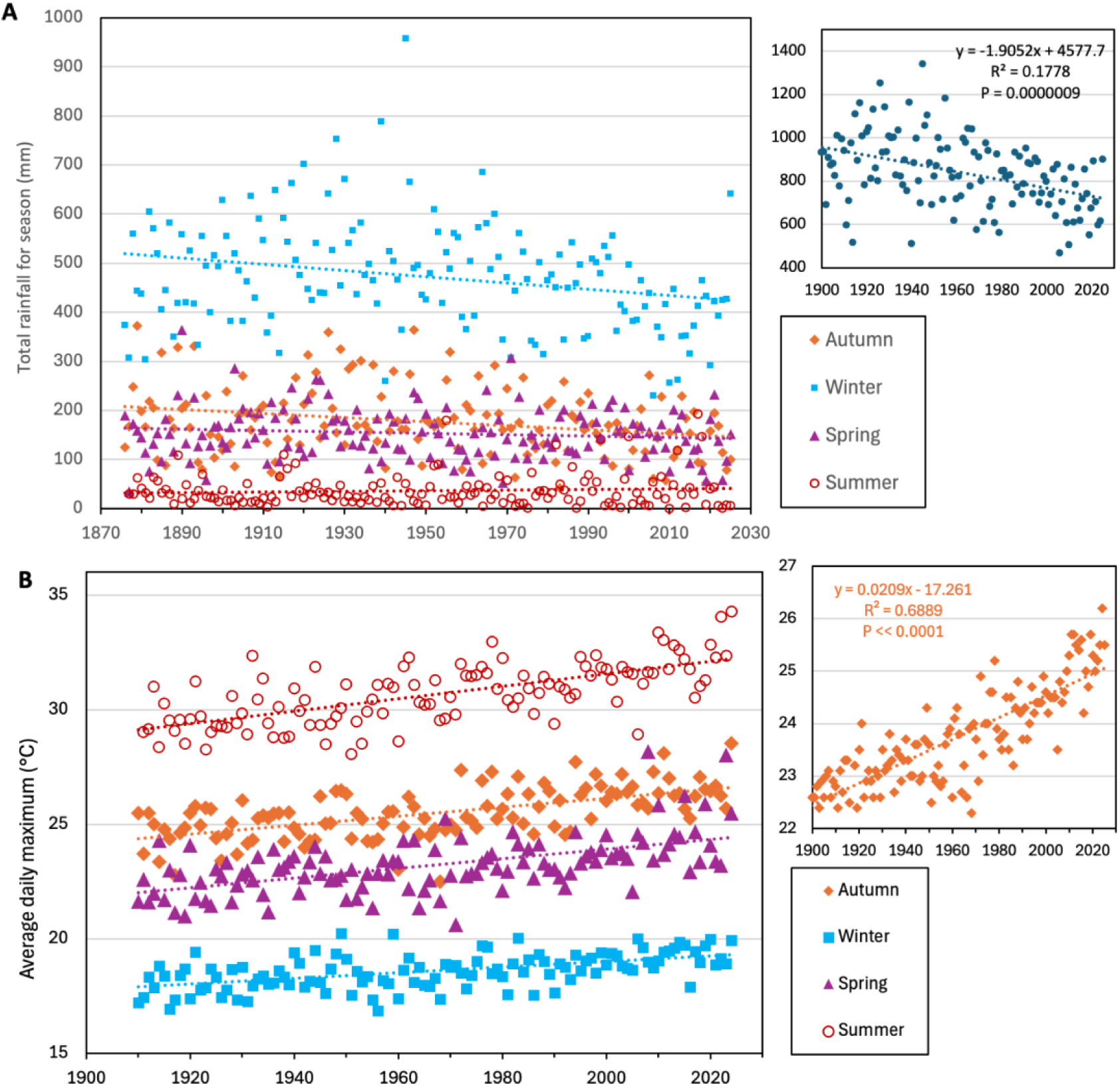
Long-term trends in rainfall (A) and temperature (B) for seasons in the Perth region (data from bom.gov.au). Insets show annual rainfall or temperature trends since 1900 and regression trend significance.

**Figure 2.**
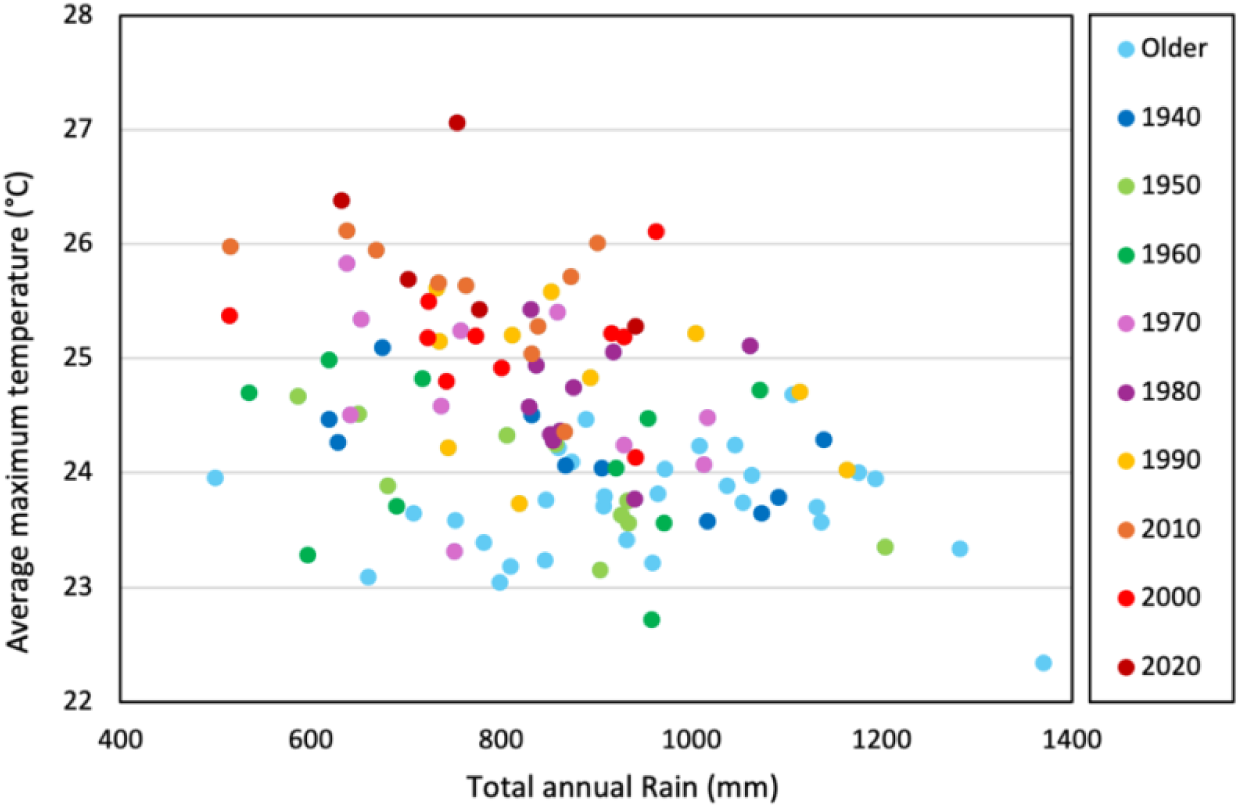
Average mean maximum temperature and total rainfall (1910-2025), showing decades becoming hotter and drier.

**Figure 3.**
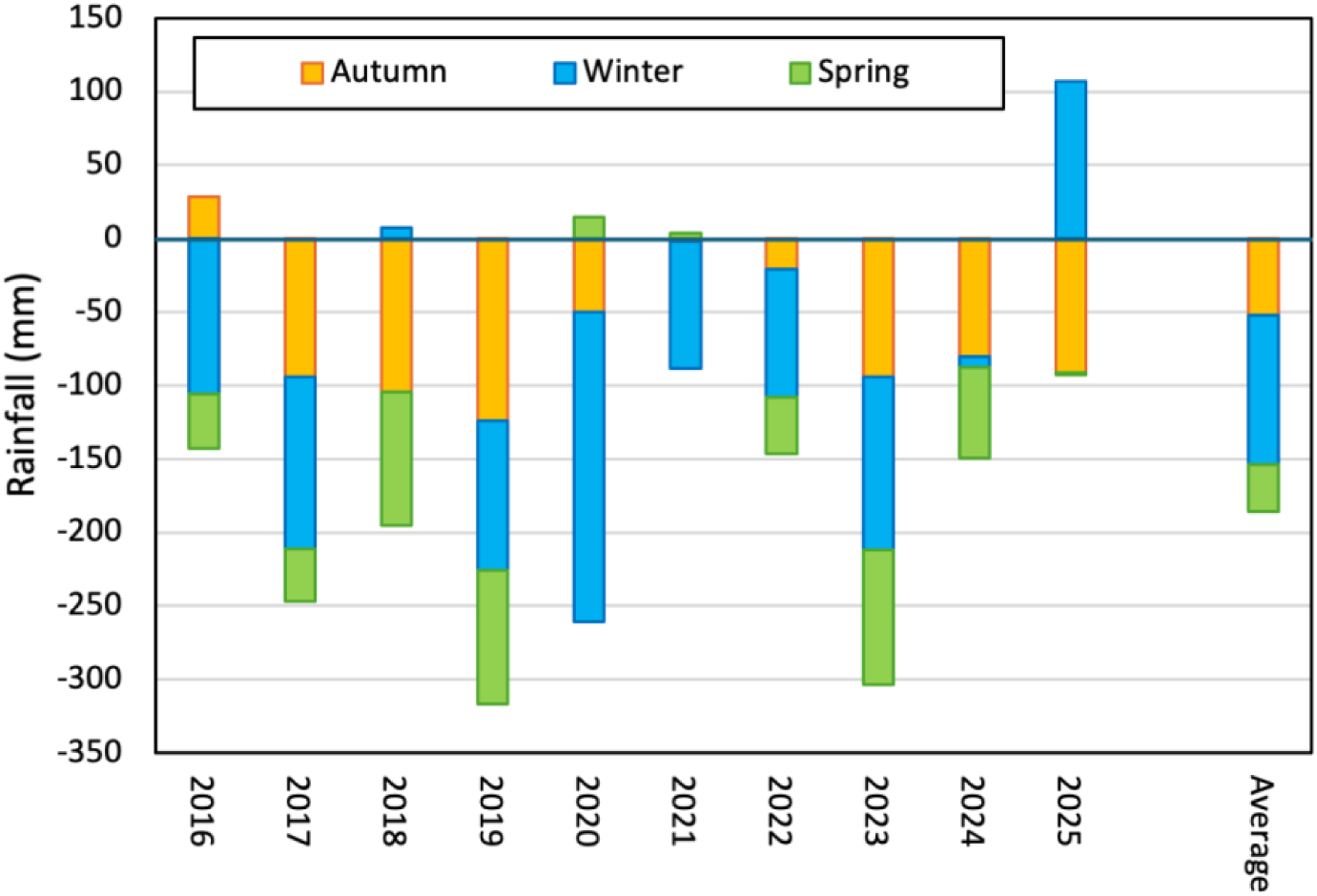
Seasonal deficits in rainfall during the study relative to the long-term average since 1906.

### 4. Orchid survey methods

Comprehensive collection of data on the abundance, flowering and pollination of orchids in Warwick Bushland began in 2016 (Brundrett 2019). Over 1000 monitoring locations for one or more species were recorded on a mobile phone using the SW Maps Android app (3-10 m accuracy) and identified using markers and georeferenced photos of surrounding features. For aggregated species, data includes both smaller and larger groups to account for density effects (Brundrett 2019). Large groups were more common in long unburnt areas and originated clonally or from seed (Brundrett 2025). Most were <10 m wide, except for *Thelymitra macrophylla* and *Leporella fimbriata* which had colonies up to 30 m across. Orchids were visited multiple times per year to record total numbers of plants, spikes, seedlings, flowers and mature capsules, and document growth stages (Brundrett 2019). Flower production per spike was used to measure the size and health of orchids and pollination to represent reproductive success (Table S2). Pollination was also converted to seed volumes per spike (yields) to compensate for major differences in capsule sizes and numbers (Table1, Fig. 4). Data from areas with recent fire was excluded for species with strong fire responses such as *L. fimbria* and *Pheladenia deformis*. Most other species had similar productivity after fire or were usually absent from recently burnt areas (Brundrett 2025).

**Figure 4.**
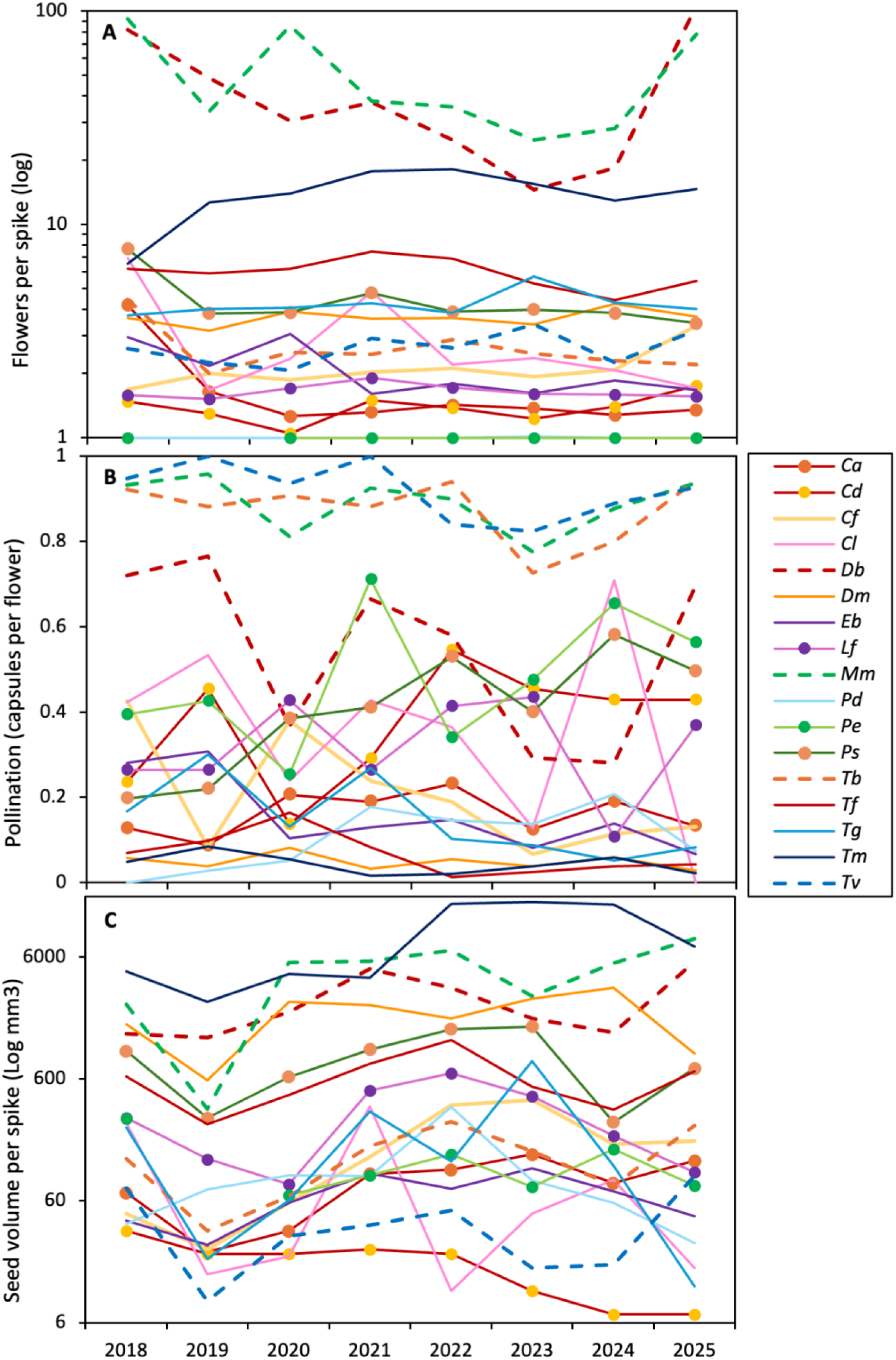
Average annual flower production (**A**), pollination (**B**) and seed production (**C**) for 17 orchids (see Table 1 for species names). Pollination is by visual deception (plain lines), sexual deception (point markers), or self-pollination (dashed lines).

### 5. Data analysis

Climatic data were amalgamated for seasons (autumn, winter and spring), bi-monthly periods and months for comparison with orchid performance, as explained in Section 2. Data used were daily average maximum temperatures for time periods (labelled as heat) and total rainfall for the same periods (rain). The climatic susceptibility of species was initially measured using graphs of seasonal rainfall or temperature data versus plant productivity, with linear regressions showing trends for species, with GLM analysis of significance. Correlation coefficients from the same climate – plant productivity comparisons were converted to indexes to quantify climate impacts on orchids, as explained in Section 3 below. These indexes were compared with orchid phenology and pollination syndromes, using regressions, correlation coefficients and GLM analysis (Section 4). Climatic responses of species were also correlated with key functional traits of orchids for reproduction and fire tolerance using previously calculated indexes in Table S6 (Brundrett 2025).

## Results and Discussion

### 1. Rainfall and temperature trends

Increasing declines in annual rainfall totalling ∼175 mm have occurred in the Perth region over the past 150 years (Fig. 1A) and the average maximum temperature increased by ∼2.6 °C over 125 years (Fig. 1B). Both trends were extremely significant (P<0.000001). Temperature and rainfall anomalies were inversely correlated with each other and became progressively more severe over 10 decades (Fig. 2). Declines in rainfall are common throughout Australia, but have increased substantially since the 1970s, especially in Western Australia (Scanlon & Doncon 2020, Verhoeven et al. 2022, Hawke et al. 2025).

Seasonal climate analyses for the study period revealed the greatest rainfall declines in winter, followed by autumn and spring, which constricted growing seasons from both ends (Figs. 1A, 3). Since 2015, several hundred mm rainfall deficits occurred in autumn on 7 out of 12 years, in winter on 9 years and in spring on 8 years, with average total deficits of −180 mm (Fig. 3). Periods of above average rainfall only occurred on single years in autumn, winter or spring (Fig. 3). There was little change to summer rainfall, when orchids were fully dormant. Increases in maximum temperature were most substantial in summer, followed by spring, autumn and winter (Fig. 1B).

### 2. Climate responses of orchids

Investigations of annual variations in pollination or flower production from numerous observations did not reveal clear trends (Fig. 4). This was caused by different seasonal trends for species, complicated by 34-fold differences in flower numbers per spike, and 26-fold capsule size variations (Fig. 5). As seen in Figure 5A, relationships between flower production and pollination were most divergent for self-pollinating orchids (*Thelymitra benthamiana, T. vulgaris*, *Disa bracteata* and *Microtis media*). Two of these (*D. bracteata* and *M. media*) also produce numerous small capsules, and strongly influence regressions (Fig. 5).

**Figure 5.**
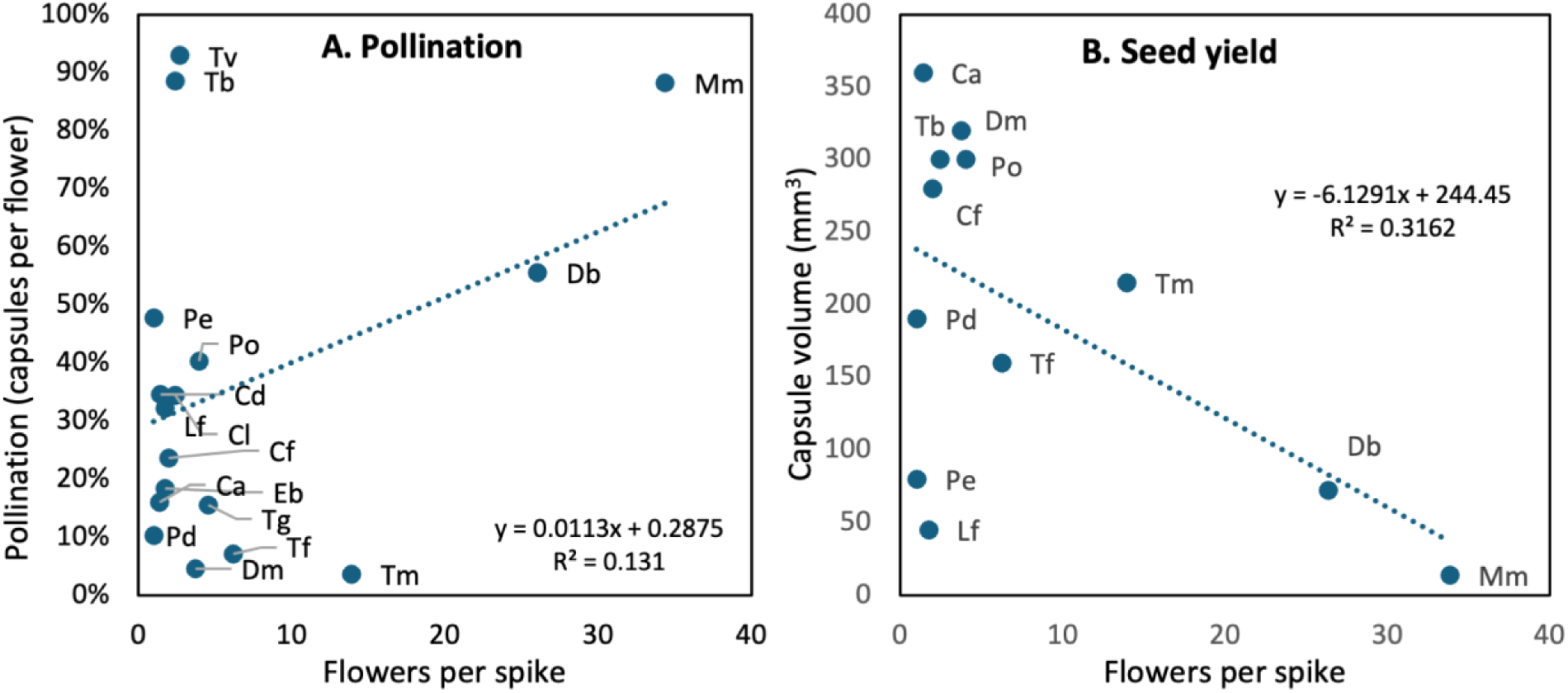
Relationship between average values for pollination (**A**) or capsule volume (**B**) relative to flower production for orchids in this study (see Table 1 for species name abbreviations).

Orchids emerged and flowered over an extend period so experienced different climatic conditions (Fig. 6). Their leaf growth periods varied over 3 months from autumn to winter, while flowering spanned seven months. Seed formation continued for 4-6 weeks after flowering for species with large capsules, but overlapped with flowering considerably for self-pollinating orchids with numerous small flowers and capsules. Thus, climatic data needed to be summarised into shorter time intervals (seasons) to account for orchid phenologies. This approach allowed single annual biological measurements (seed, flower production, etc.) to be contrasted with continuously measured environmental data (rain, temperature, etc). A second major benefit was that contrasting species with climate individually compensated for major differences in flowering and seed production without data transformation (Fig. 5). A third benefit was that separate comparisons revealed climatic factors or productivity measures with both positive and negative effects on the same orchid, that otherwise could negate each other (see Section 4).

**Figure 6.**
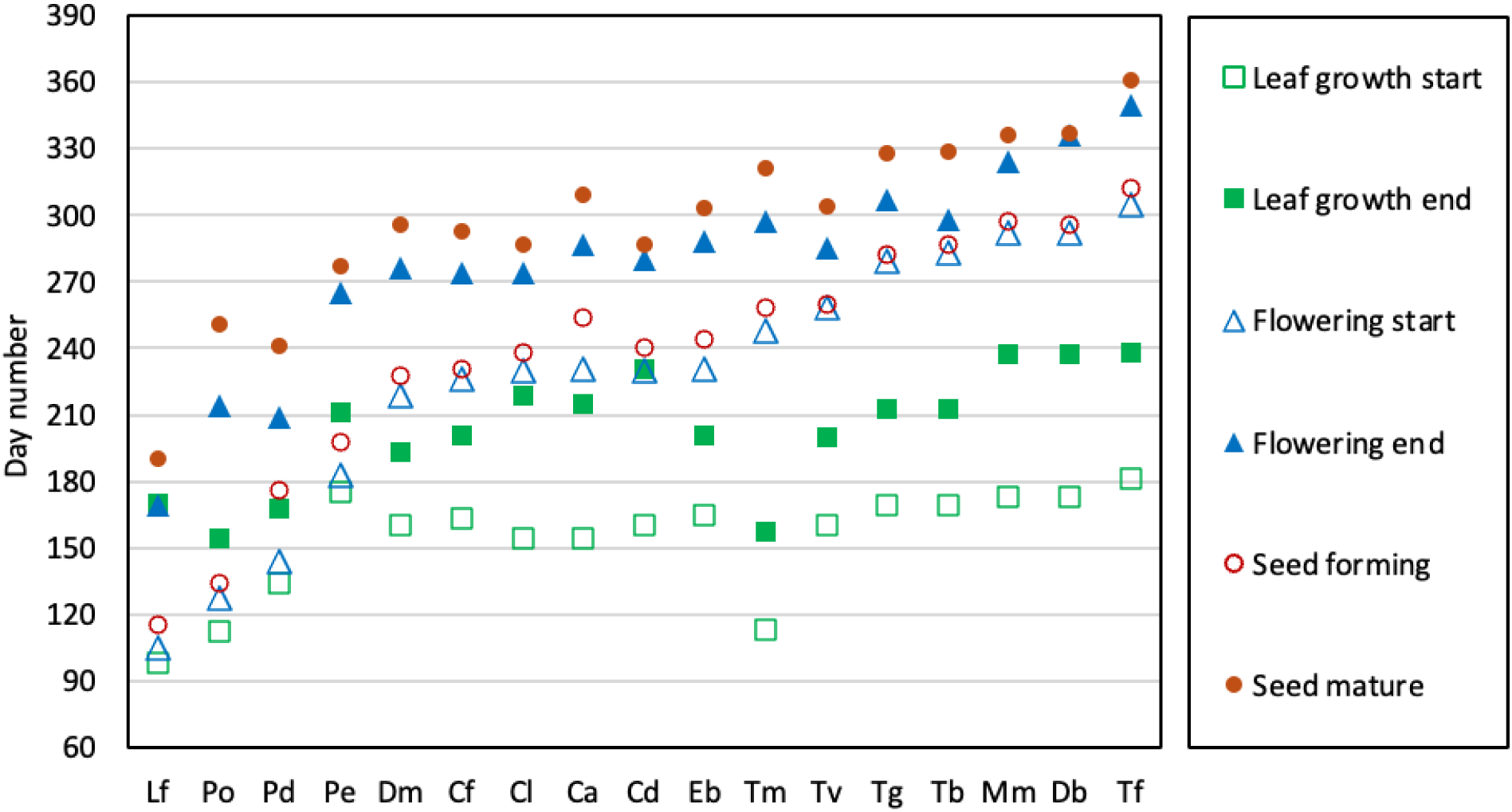
Comparative phenology of orchids showing average start or end dates for leaf growth, flowering and seed production (see Table 1 for species names).

Orchids in this study (and the entire SWAFR) were frequently exposed to major climatic anomalies (Fig. 3). These rainfall and temperature fluctuations and orchid productivity data were combined in climate response graphs that summarised orchid responses (Figs. 7-10). Separate graphs contrasted plant productivity against autumn, winter and spring rainfall totals or average maximum temperatures (heat). Comparisons of annual rainfall totals for the current and previous year were less informative (not presented). Annual growing season transition dates determined by rainfall patterns were also used in climate analyses. Figures 7 to 10 include 24 graphs with relatively strong trends (R^2^ values > 0.2). These include 102 climate - orchid response comparisons, of which 26% were significant (P <0.01), resulting in at last one significant trend for 14 of the 17 species. Many graphs show marginally significant but very similar trends for multiple species, demonstrating response consistency, while others show species with divergent responses to climate (e.g. Fig. 7D, F).

**Figure 7.**
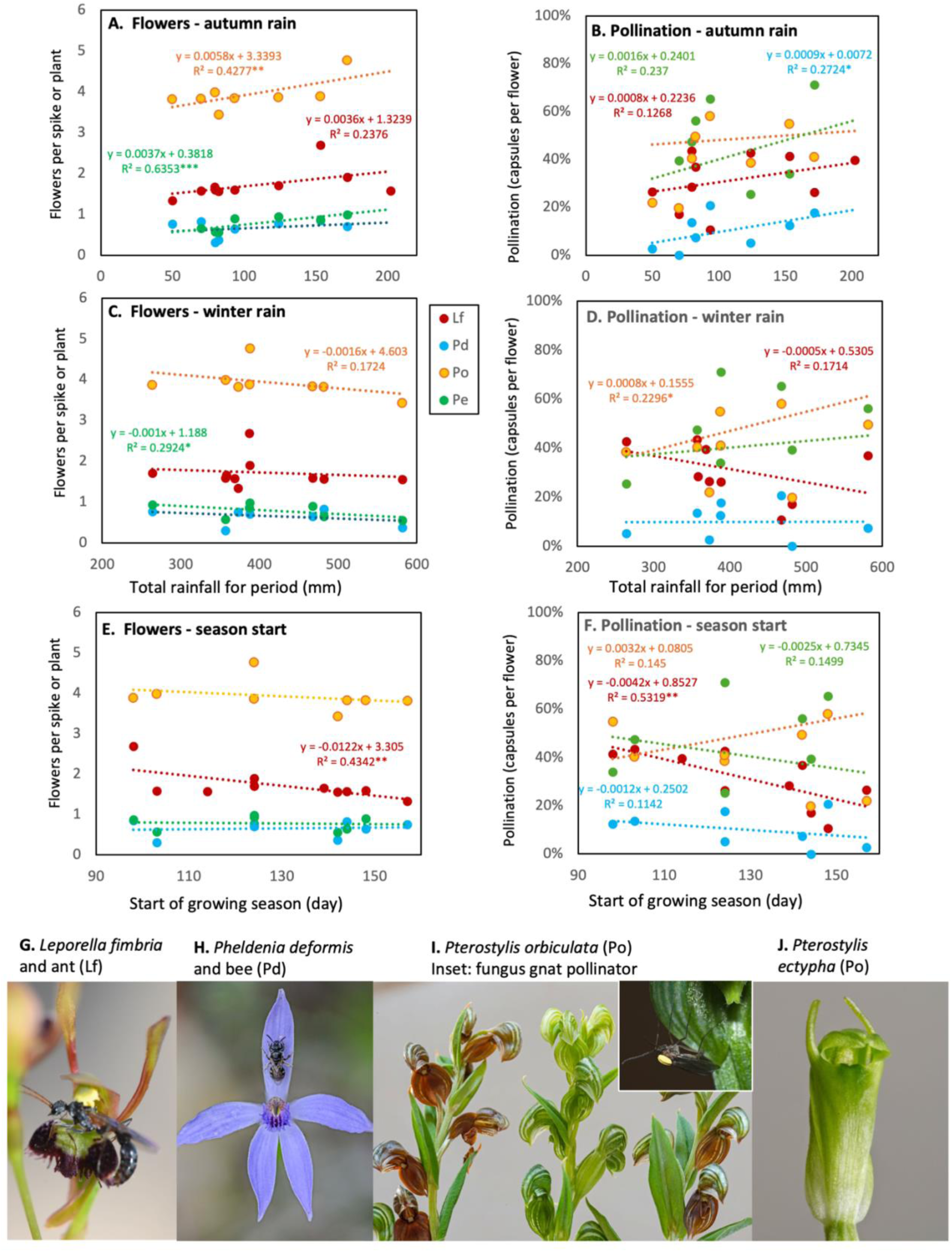
Effects of seasonal rainfall totals on four autumn or winter flowering orchids. Climate data are for autumn rain (**A, B**), winter rain (**C, D**), or start of the growing season (**E, F**). See Table 1 for species names (* = P< 0.1, ** = P< 0.05, *** = P< 0.01). Photos of orchids and pollinators (**G-J**).

Overall, seasonal trends were strongest for autumn climate (66% were for autumn rain or season start day), followed by spring rain. Winter rain was adequate every year and summer rain was sparse on all but two years. Regarding impact types, there was an even split between pollination (48%) and flower production (52%) for significant trends. The former measured combined impacts on insects and plants, while the latter only concerns plant responses. Climate responses are organised into four groups below that separate species by flowering phenology and pollination syndromes.

#### A. Flowering in autumn or winter

This group includes three sexually deceptive orchids (*L. fimbria, Pterostylis orbiculata* and *Pterostylis ectypha*) and a winter-flowering visually deceptive orchid (*Pheladenia deformis*). *Leporella fimbria* (hare orchid – Fig. 7G) was first to flower in autumn and early winter. This orchid has fire-promoted flowering and is pollinated by male bull ants (*Myrmecia* spp.) that fly to find mates following wet autumn weather (Peakall 1989, Brundrett 2019, Kuiter 2025, www.youtube.com/watch?v=D1FTrJaEFhE&t=68s). Flower production varied substantially between locations, with most plants remining sterile (0-15% of plants in a location flower). *Leporella fimbria* flowering was significantly impacted by autumn, but not winter drought, while pollination was only weakly affected by rain (Fig. 7A-D). Both flowering and pollination was substantially impacted by delayed growing seasons (Fig. 7E, F).

*Pterostylis orbiculata* (banded greenhood, also known as *P. sanguinea* – Fig. 7I) has shoots emerging in autumn and flowers in winter (Fig. 6). Its flowering was reduced by autumn drought (Fig. 7C) and winter rain was weakly detrimental (Fig. 7C), probably due to loss of flowers to fungal infections (Brundrett 2019). Pollination increased with winter rain and earlier season starts (Fig. 7B-F). Thus, pollination by fungus gnats (Phillips et al. 2013, www.youtube.com/watch?v=9JVNQzovcGQ) is relatively tolerant of wet cool conditions.

*Pterostylis ectypha* (snail orchid – Fig. 7J), is a small, short-lived colonial, single-flowered, disturbance tolerant, orchid killed by fire (Brundrett 2025). This *Pterostylis* species is also fungus gnat pollinated, flowers in late winter to early spring and benefits substantially from increased autumn but not winter rain (Fig. 7A-D). Both *Pterostylis* species have relatively efficient pollination compared to visually deceptive and other sexually deceptive species (Brundrett 2019).

*Pheladenia deformis* (bluebeard orchid – Fig. 7H) is a small, short-lived species spreading by seed that flowers in mid-winter (Brundrett 2025). This orchid has single flowers, with large annual variations in flowering and very low seed production, except after fire. Its pollination, presumably by bees (Fig. 7H), benefits from autumn, but not winter rain, and flowering was not affected by either (Fig. 7A-D). Growth, flowering and seed production occurs primarily in winter, so drought stress is normally avoided. However, flowering the previous year can reduce seed production due to reproductive costs (Brundrett 2025). Both *P. ectypha* and *P. deformis* consistently have singe flowers so are represented by the proportion of plants with a flower (Fig. 7).

#### B. Spring flowering visually deceptive orchids

These orchids flower in sequence over an extended period from late winter to early spring for *Caladenia flava* (cowslip orchid – Fig. 8G), *C. latifolia* (pink fairy orchid - Fig. 8H) and *Diuris magnifica* (pansy orchid - Fig. 8I), followed by *Elythranthera brunonis* (enamel orchid - Fig. 8J)*, T. macrophylla* (blue sun orchid - Fig. 8M) and *T. graminea* (slender blue sun orchid - Fig. 8L) in mid spring, then *T. fuscolutea* (chestnut orchid - Fig. LK) in late spring (Fig. 6). Flower production for three species benefitted substantially from autumn rain (Fig. 8A), with the strongest benefits for *T. macrophylla* and *C. latifolia*. Flowering responses to winter rain, were negative and weak, expect for *T. macrophylla*. Pollination had limited responses (Fig. 8C-E), except for positive effects of autumn rain on *E. brunonis* and winter rain on *C. latifolia*. Growing season length (Fig. 8F) and spring rainfall were less important, especially for late flowering sun orchids which are relatively drought tolerant. *Thelymitra macrophylla* is an outlier in this group with flower production strongly promoted by autumn rain (Fig. 8A) and longer seasons (Fig. 8F), which is due to very early shoot emergence (Fig. 6). In general, benefits of increased rainfall for visually deceptive orchids, seem to be offset by reduced insect activity due to lower temperatures, especially for sun orchids (Fig. 8E).

**Figure 8.**
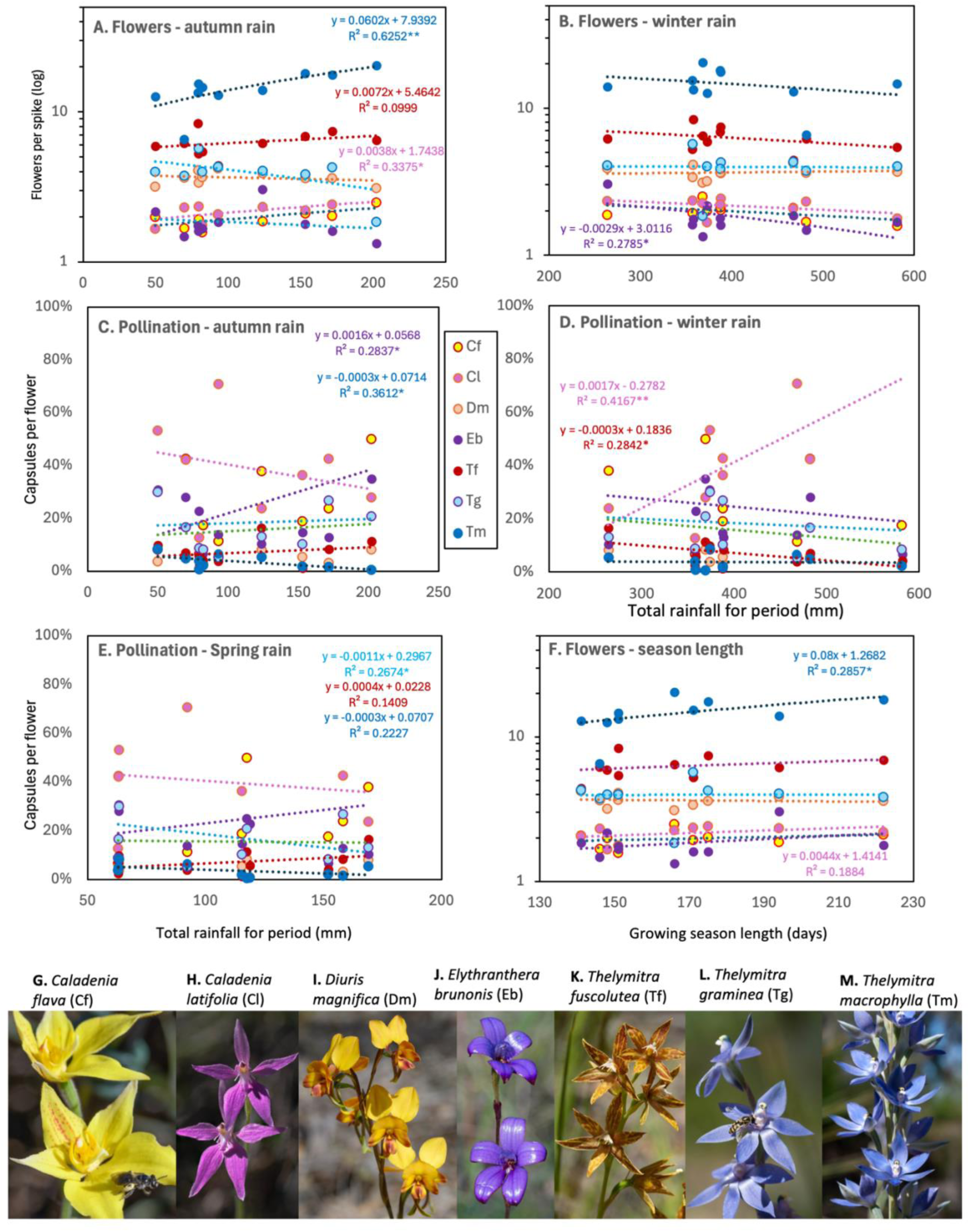
Effects on flowering (**A, B, F**) or pollination (**C, D, E**) of seven spring flowering visually deceptive orchids relative to seasonal rainfall (**A-E**) or growing season length (**F**). See Table 1 for species names (* = P< 0.1, ** = P< 0.05). Photos of orchids and pollinators (**G-M**).

#### C. Spring flowering sexually deceptive orchids

Two spider orchids (*Caladenia arenicola,* carousel spider orchid- Fig. 9G and *C. discoidea*, dancing orchid – Fig. 9H) have similar phenology but differ substantially in size and resilience (the former is a larger plant with stable numbers, while the latter became less common). Both are pollinated by male thynnine wasps attracted by pheromones (Fig. 9GH, www.youtube.com/channel/UCw3FAqI4-PXG8MxKhFYCwNQ). *Caladenia arenicola* flowered more with increased autumn rain (Fig. 9A, B), and yielded more seeds with longer growing seasons (Fig. 9E, F). Spring rain was beneficial for *C. arenicola,* but not *C. discoidea*, where autumn and winter rain had contrasting effects (Fig. 9C, D). Wasps that pollinate sexually deceptive *Caladenia* species generally seem to have broader temperature tolerances than pollinators of visually deceptive species (mostly indigenous bees).

**Figure 9.**
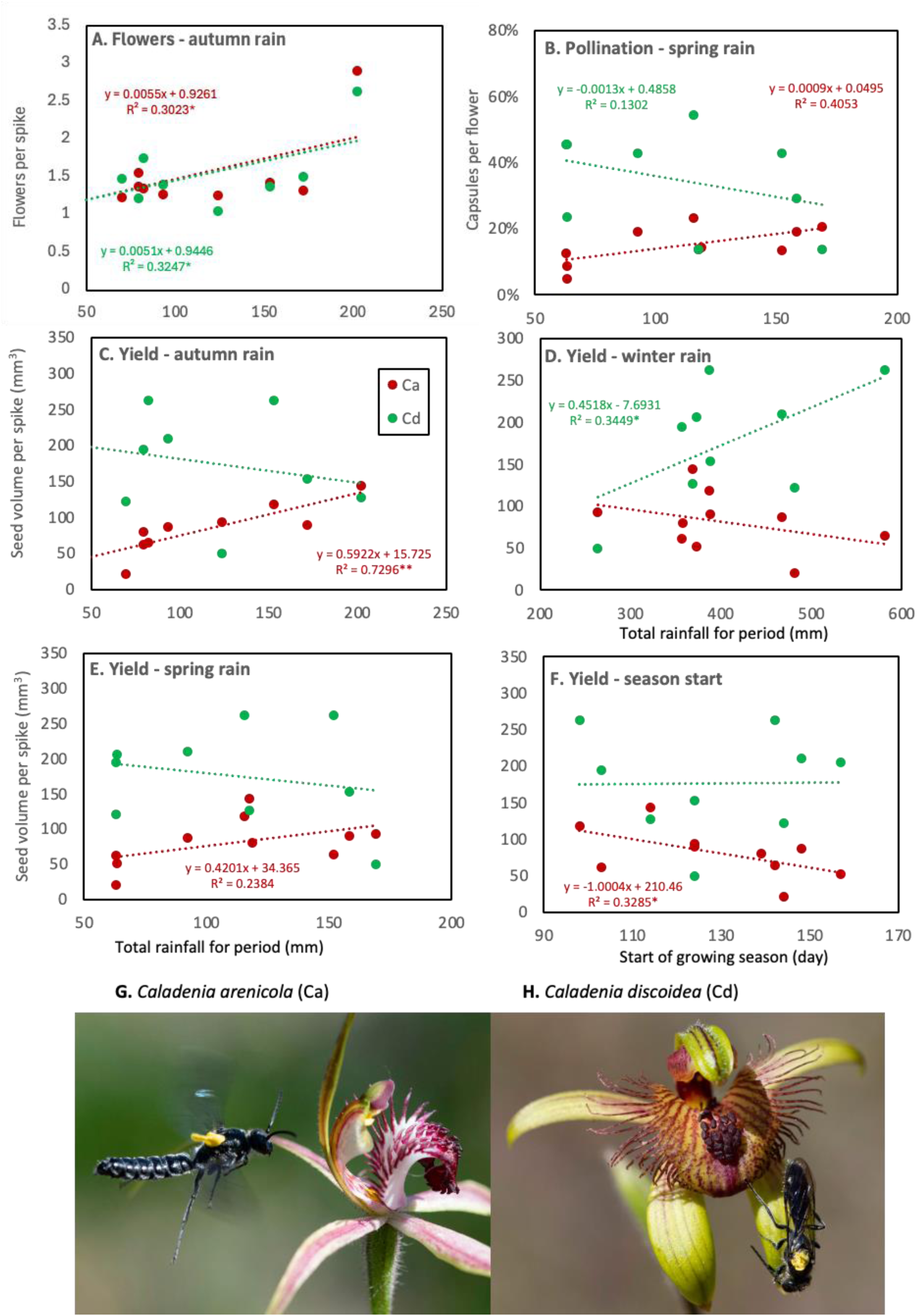
Effects of seasonal rainfall on flowering (**A, C, E**) or pollination (**B, D, F**) of spring flowering sexually deceptive orchids (* = P< 0.1, ** = P< 0.05). Photos show attraction of thynnine wasps carrying pollinia (**G-H**).

#### D. Self-pollinating orchids

Three of these species flower in late spring (Fig. 6), so yield was affected by drought when flowers and seeds are forming (Fig 10). These include two short-lived, disturbance tolerant, weedy orchids that reproduce only by seed and are killed by fire. These are *Microtis media* (mignonette orchid - Fig. 10H) and *Disa bracteata* (South African orchid - Fig. 10G). *Thelymitra benthamiana* (leopard orchid - Fig. 10I) is a sun orchid flowering in late spring, that prefers undisturbed and long unburnt habitats (Brundrett 2025). The remaining species, *Thelymitra vulgaris,* (small blue sun orchid – Fig. J) produces small flowers in early spring. Both sun orchids (*T. vulgata* and *T. benthamiana*), have flowers that only open a few times on warm days. Their flowering is resilient to rainfall variations in all seasons (Fig. 10A-C). Both species have weak climatic responses for pollination, as they lack insect dependency (Fig. 10D), but autumn rain benefited *T. benthamiana* (Fig. 10A). *Disa bracteata*, which is invasive from South Africa, had strong positive flowering responses to autumn and spring rain, but *M. media*, which is indigenous, was more resilient (Fig. 10A-C). Flowering and seed production for *M. media* were also impacted by earlier growing season end dates (Fig 10E, F). Both species have numerous semi-indeterminate very small flowers, with production determined by climate and plant vigour (Brundrett 2019).

**Figure 10.**
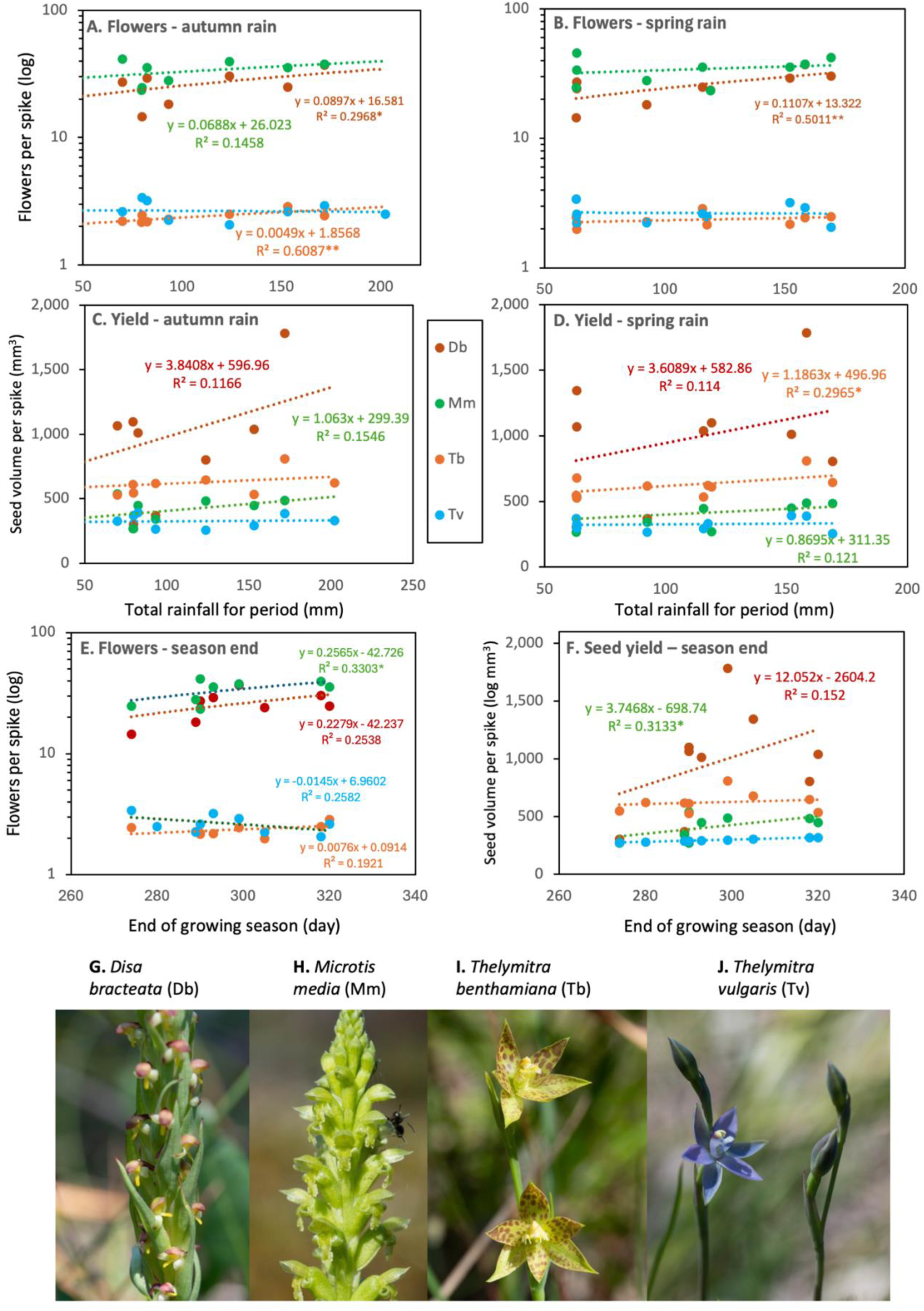
Effects of seasonal rainfall on flowering (**A –C, E**) or pollination (**D, F**) of four spring flowering self-pollinating orchids (* = P < 0.1, ** = P <0.05). Photos of orchids (**G-J**).

### 3. Overall orchid responses to climate

A new framework for quantifying orchid-climate relationships of species was developed using correlations between rainfall or maximum temperature and flower or seed production for the same years (see Table 2). This resulted in four tables of correlation coefficients for all orchid species (Tables 2, S3). Only correlations between orchids and climate were utilised, as orchid-orchid and climate-climate correlations were less informative and repeated across multiple tables (Tables 2C, S4). Species responses summarised in Figure 11 classified correlations into strong negative, negative, neutral, positive or strong positive categories using a consistent scale (>−0.3, >−0.1, −0.1 to 0.1, <0.1, <0.3). These were also compiled over three different time scales (monthly, bi-monthly and seasonally) to determine which corresponds best to orchid responses. Climatic data were excluded for summer months when orchids were fully dormant, rain was sparse and erratic, and temperatures were consistently warm or hot (Table S2).

**Figure 11.**
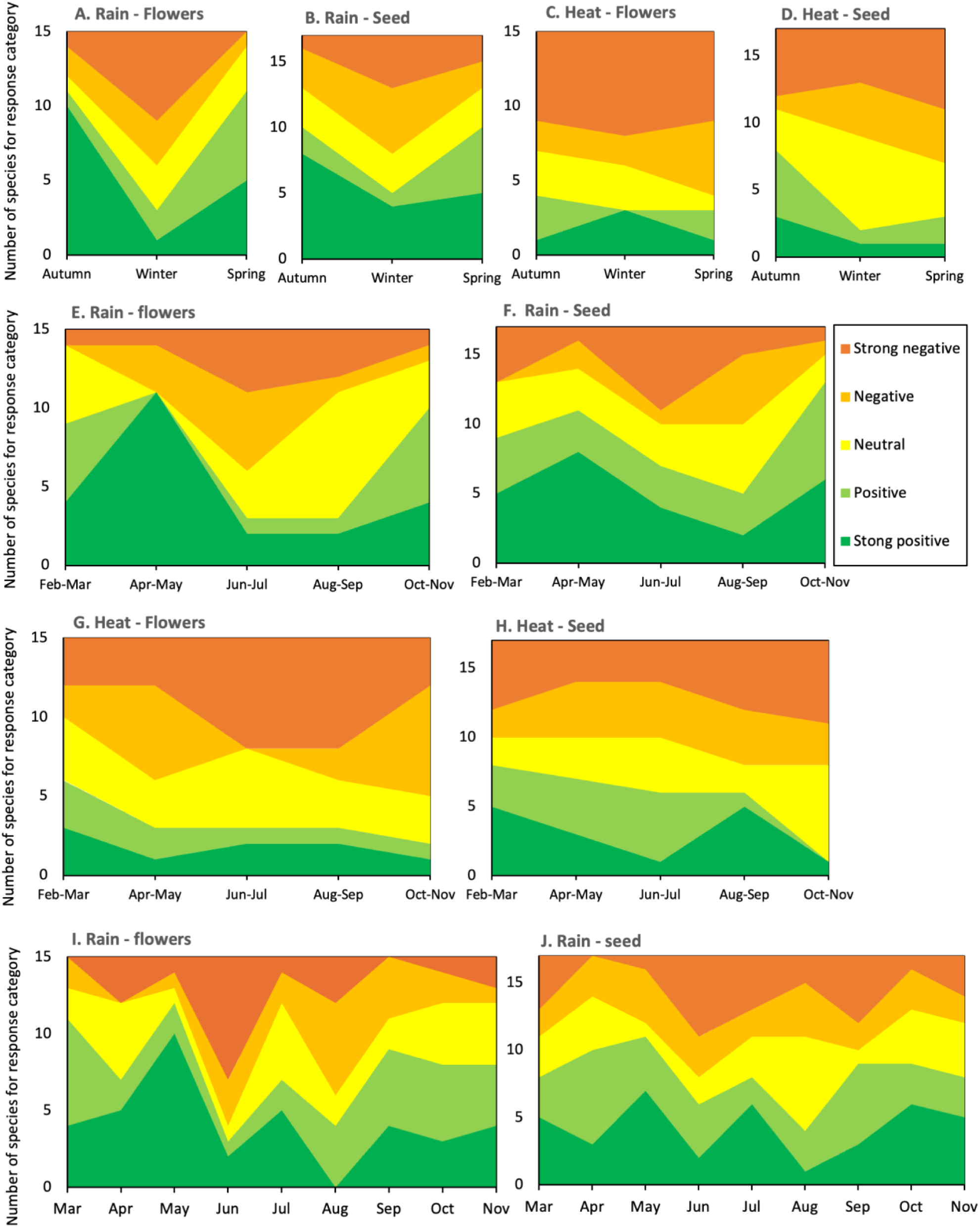
Climate response categories for orchid species based on correlations between rainfall (**A, B, E, F**) or temperature (**C, D, G, H**) and flower production per spike (**A, C, E, G**), or seed volume per spike (**B, D, F, H**). Climatic data are compiled using conventional seasons (3 monthly – **A-D**), approximated Noongar seasons (2 monthly – **E-H**), or monthly records (**I, J**).

**Table 2.** Use of correlations to calcuate climate indexes for orchid species.

| A. Climate data |  | Total rainfall |  |  | Average maximum temperature |  |  |  | Growing season |  |  |
| --- | --- | --- | --- | --- | --- | --- | --- | --- | --- | --- | --- |
| Year | Summe | Autumn | Winter | Spring | Summer | Autumn | Winter | Spring | Start of | End of |  |
| 2016 | 30.7 | 202 | 368 | 118 | 31.2 | 25.1 | 18.0 | 22.7 | 114 | 280 |  |
| 2017 | 173 | 79 | 358 | 119 | 29.8 | 25.9 | 19.5 | 24.4 | 139 | 290 |  |
| 2018 | 118.7 | 70 | 482 | 63 | 30.1 | 26.6 | 18.9 | 23.1 | 144 | 290 |  |
| 2019 | 6.6 | 50 | 373 | 63 | 30.5 | 26.0 | 19.8 | 25.3 | 157 | 305 |  |
| 2020 | 30.6 | 124 | 264 | 169 | 32.0 | 26.0 | 19.9 | 23.9 | 124 | 318 |  |
| 2021 | 49.6 | 172 | 388 | 158 | 31.6 | 26.4 | 18.9 | 23.0 | 124 | 299 |  |
| 2022 | 2.4 | 153 | 387 | 116 | 33.3 | 26.1 | 19.2 | 22.9 | 98 | 320 |  |
| 2023 | 0 | 80 | 357 | 63 | 31.4 | 25.3 | 18.8 | 26.3 | 103 | 274 |  |
| 2024 | 1.9 | 93 | 467 | 92 | 32.6 | 27.8 | 19.9 | 24.5 | 148 | 289 |  |
| 2025 | 5.05 | 82 | 581 | 152 | 31.6 | 27.6 | 19.2 | 23.6 | 142 | 293 |  |
|  |  | Period total (mm) |  |  |  | Period average (°C) |  |  | Transition day no |  |  |
| B. Orchid productivity data (flowers per spike) |  |  |  |  |  |  |  |  |  |  |  |
| Year | Ca | Cf | Db | Dm | Eb | Lf | Tf | Tm | See Table 1 for names |  |  |
| 2016 | 2.90 | 2.50 |  | 3.1 | 1.3 | 1.6 | 6.5 | 20.5 | Data from Table S1 |  |  |
| 2017 | 1.56 |  | 23.7 | 4.1 | 1.8 | 1.7 | 8.4 | 13.4 |  |  |  |
| 2018 | 1.23 | 1.68 | 27.3 | 3.6 | 1.5 | 1.6 | 6.2 | 6.6 |  |  |  |
| 2019 | 1.64 | 2.00 | 24.3 | 3.2 | 2.2 | 1.3 | 5.9 | 12.6 |  |  |  |
| 2020 | 1.26 | 1.87 | 30.4 | 3.9 | 3.1 | 1.7 | 6.2 | 14.0 |  |  |  |
| 2021 | 1.32 | 2.03 | 37.3 | 3.6 | 1.6 | 1.9 | 7.4 | 17.7 |  |  |  |
| 2022 | 1.42 | 2.10 | 24.9 | 3.6 | 1.8 | 2.7 | 6.9 | 18.1 |  |  |  |
| 2023 | 1.37 | 1.93 | 14.6 | 3.4 | 1.6 | 1.6 | 5.3 | 15.4 |  |  |  |
| 2024 | 1.27 | 2.08 | 18.4 | 4.2 | 1.8 | 1.6 | 4.4 | 12.9 |  |  |  |
| 2025 | 1.35 | 1.58 | 29.3 | 3.7 | 1.7 | 1.6 | 5.4 | 14.7 |  |  |  |
| C. Correlations between seasonal rainfall and orchid flowering over 9-10 years |  |  |  |  |  |  |  | Data from Table S2 |  |  |  |
| Orchid | Autumn rain | Winter rain | Spring rain | Ca | Cf | Db | Dm | Eb | Lf | Tf | Tm |
| Autumn rain | 1 | Rain-rain correlations (not used) |  |  |  |  |  |  |  |  |  |
| Winter rain | -0.289 | 1 |  |  |  |  |  |  |  |  |  |
| Spring rain | 0.535 | -0.109 | 1 |  |  |  |  |  |  |  |  |
| Ca | 0.550 | -0.192 | -0.018 | 1 | Orchid-orchid correlations (not used) |  |  |  |  |  |  |
| Cf | 0.699 | -0.476 | -0.027 | 0.770 | 1 |  |  |  |  |  |  |
| Db | 0.545 | 0.025 | 0.708 | -0.209 | -0.224 | 1 |  |  |  |  |  |
| Dm | -0.227 | 0.118 | 0.292 | -0.580 | -0.352 | -0.040 | 1 |  |  |  |  |
| Eb | -0.171 | -0.528 | 0.346 | -0.347 | -0.199 | 0.162 | 0.281 | 1 |  |  |  |
| Lf | 0.487 | -0.138 | 0.285 | -0.154 | 0.183 | 0.169 | 0.133 | -0.030 | 1 |  |  |
| Tf | 0.316 | -0.375 | 0.339 | 0.157 | 0.233 | 0.475 | 0.025 | -0.092 | 0.354 | 1 |  |
| Tm | 0.791 | -0.293 | 0.452 | 0.566 | 0.689 | 0.135 | -0.340 | -0.149 | 0.415 | 0.206 | 1 |
| Data used below |  |  |  |  |  |  |  |  |  |  |  |
| D. Index calculations from rain and flowering correlations |  |  |  |  |  |  |  |  |  |  |  |
|  | Main correlation coefficients |  |  | Correlations for all seasons |  | Min + Max | Max - Min | CSI + CRI |  |  |  |
| Orchid | Autumn rain | Winter rain | Spring rain | Max | Min | CRI | CSI | CI | CRI rank | CSI rank | CI rank |
| Ca | 0.55 | -0.19 | -0.02 | 0.55 | -0.19 | 0.36 | 0.74 | 1.10 | 6 | 5 | 5 |
| Cf | 0.70 | -0.48 | -0.03 | 0.70 | -0.48 | 0.22 | 1.18 | 1.40 | 4 | 8 | 6 |
| Db | 0.54 | 0.03 | 0.71 | 0.71 | 0.03 | 0.73 | 0.68 | 1.42 | 8 | 3 | 7 |
| Dm | -0.23 | 0.12 | 0.29 | 0.29 | -0.23 | 0.07 | 0.52 | 0.58 | 3 | 1 | 1 |
| Eb | -0.17 | -0.53 | 0.35 | 0.35 | -0.53 | -0.18 | 0.87 | 0.69 | 1 | 6 | 3 |
| Lf | 0.49 | -0.14 | 0.29 | 0.49 | -0.14 | 0.35 | 0.63 | 0.97 | 5 | 2 | 4 |
| Tf | 0.32 | -0.38 | 0.34 | 0.34 | -0.38 | -0.04 | 0.71 | 0.68 | 2 | 4 | 2 |
| Tm | 0.79 | -0.29 | 0.45 | 0.79 | -0.29 | 0.50 | 1.08 | 1.58 | 7 | 7 | 8 |

Figure 11 reveals complex orchid responses to rainfall and temperature. Most species benefitted from years with greater autumn and/or spring rainfall, while increased winter rainfall was rarely beneficial and could be detrimental (Fig. 11A, B, E, F). Years with higher temperatures were detrimental or neutral for most orchids (Fig, 11C, D, G, H), but this may be an indirect effect since temperature was correlated with rainfall (Fig. 2). Some orchids received substantial benefits from rainfall in autumn before they were growing (Fig. 11), as discussed below (Section 7).

Three different climate recording periods are compared in Figure 11. The first two were traditional three-month seasons (Fig. 11A-D), and the approximated six-season indigenous calendar (Wadjuk Noongar). These both produced similar trends, but Noongar seasons revealed more complexity during Bunuru, or early autumn (Fig. 6E-H). Noongar seasons are based on the phenology of animals and plants as well as weather, so their timing varies annually (australiassouthwest.com/six-seasons-of-the-south-west). For practical reasons, they are usually generalised using climate data from the closest two months (April-May, June-July, etc.). Response categories based on individual months had similar overall trends to seasons, but were less consistent (Fig. 11I, J).

Figure 12 compares autumn, winter and spring climate correlations with orchid productivity for orchid species arranged in flowering sequence. Flowering and seed production benefited from increased autumn and spring rainfall and were unaffected or weakly supressed by increased winter rainfall. Specifically, 10 orchids responded strongly to increased autumn rain, three late flowering species benefited from spring rain, and one preferred less rain when flowering (*T. macrophylla*). Three orchid species were strongly adversely affected by increased rainfall at different times of the year (Fig. 12). Responses to heat were not as well correlated with phenology, but three species showed strong positive flowering responses, and 10 had negative responses, especially in spring. Except for *T. macrophylla*, seed production was adversely impacted by heat, especially for late-flowering orchids.

**Figure 12.**
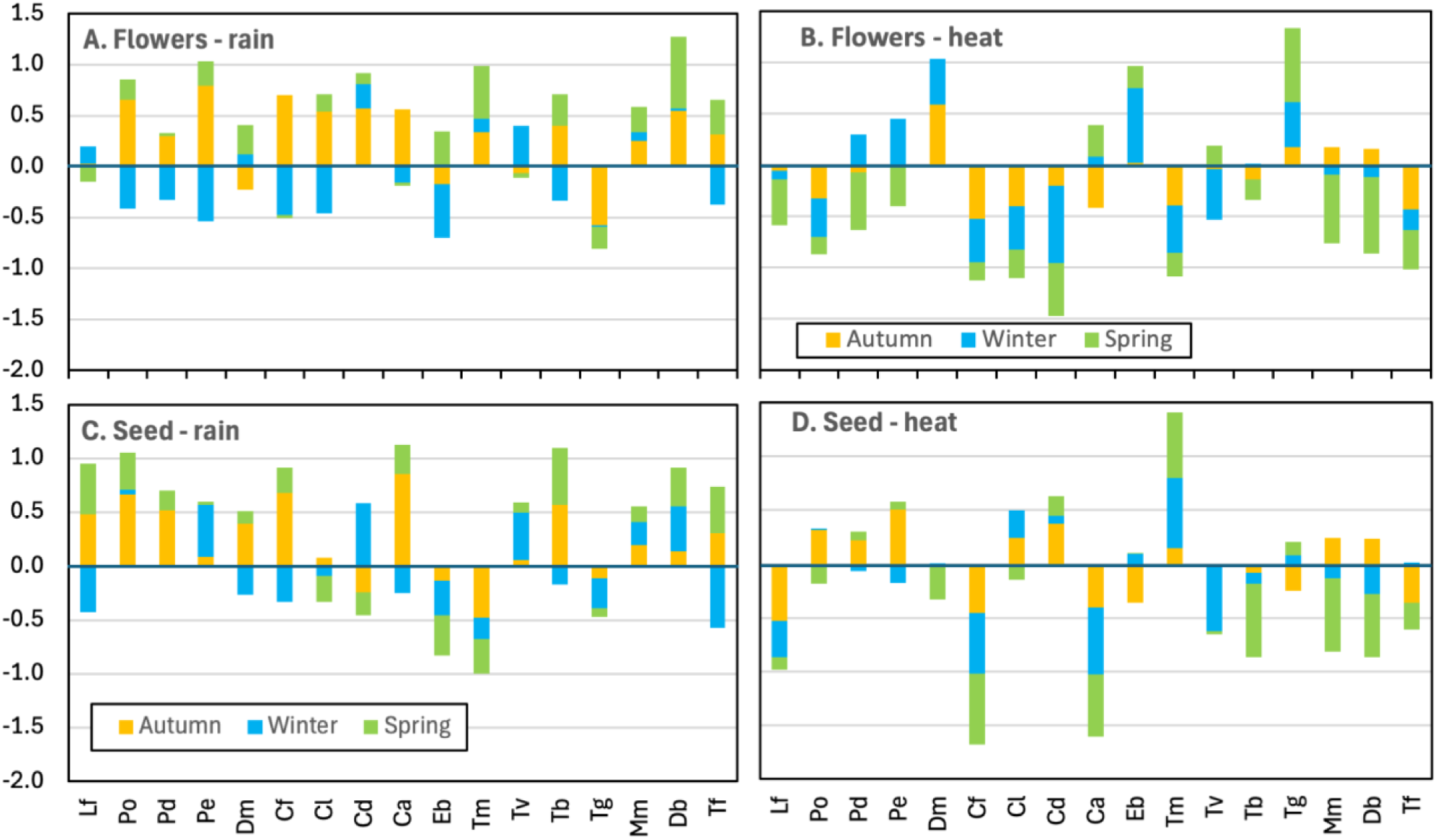
Seasonal correlation coefficients for rain (**A, C**) or heat (**B, D**), relative to flower (**A, B**) or seed production per spike (**C, D**). Orchids are arranged in flowering sequence (see Table 1 for species names).

### 4. Climate Sensitivity Indexes

Due to the complexity of relationships revealed in Figures 7 to 12, a more consistent approach was required to assign orchids to response categories. After comparing alternative workflows, the best result was achieved by converting correlation coefficients into climate indexes, as explained in Box 1 and Table 2. These correlation coefficients were produced from orchid productivity and climate data, as explained above. Indexes that were based on maximum and minimum correlation values across all seasons produced the best results (Box 1). Their sum forms the Climate Response Index (CRI), which is equal to twice the median (Table 2). Subtracting the minimum from the maximum correlation value provided the Climate Sensitivity Index (CSI), showing the span of all values. Examples of index values are presented in Table 3. and compared in Figure 13, which shows that the CSI was always positive and represented the magnitude of responses, while the CRI revealed the direction of climate responses (positive or negative), so was more informative. The CI index (CRI and CSI combined) was less useful because it was dominated by the CRI (Fig. 13). Consequently, CRIs were used for most analyses, but CSIs were combined with other data for climate sensitivity ordinations (see below).

**Figure 13.**
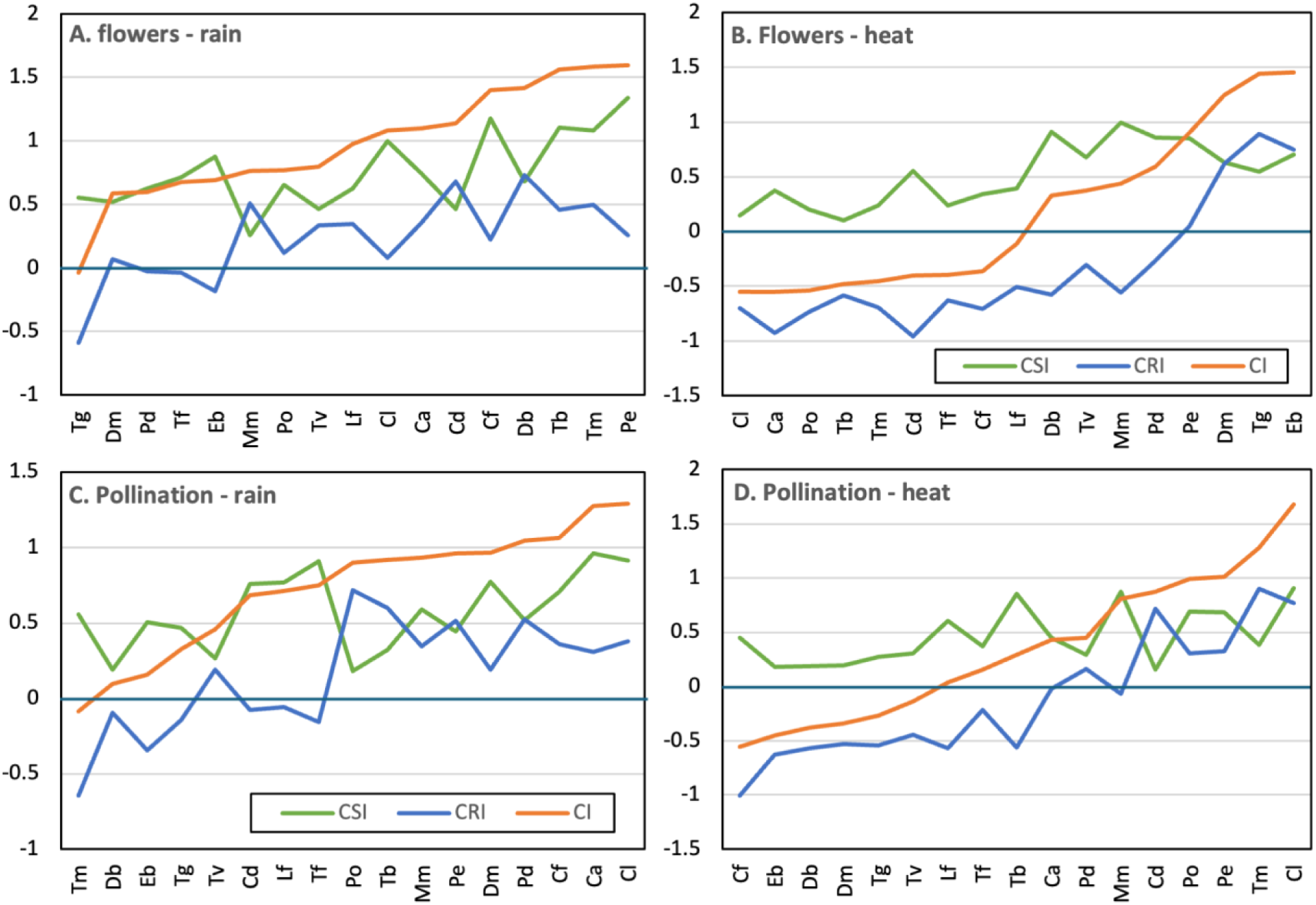
Climate response (CRI), sensitivity (CSI) and combined Indexes (CI) for rainfall or temperature effects on flower production or pollination. Orchids are sorted by CI values (see Table 1 for species names).

**Table 3.** Summary of all indexes with ranks for species.

| A. CRI Table |  |  |  |  |  |  |  |  |  | Rain rank |
| --- | --- | --- | --- | --- | --- | --- | --- | --- | --- | --- |
| Orchid | R-F CRI | R-P CRI | R-SV CRI | H-F CRI | H-P CRI | H-SV CRI | Rain CRI | Heat CRI | Total CRI |  |
| Ca | 0.36 | 0.31 | -0.71 | 0.92 | -0.02 | 0.73 | -0.35 | 1.66 | 1.31 | 1 |
| Cd | 0.68 | -0.08 | -0.68 | 0.96 | 0.72 | -0.45 | 0.00 | 0.51 | 0.51 | 5 |
| Cf | 0.22 | 0.36 | -0.36 | 0.71 | -1.00 | 1.10 | -0.14 | 1.81 | 1.67 | 4 |
| Cl | 0.08 | 0.38 | 0.34 | 0.70 | 0.77 | -0.77 | 0.42 | -0.07 | 0.35 | 8 |
| Db | 0.73 | -0.09 | 0.58 | 0.58 | -0.57 | 0.60 | 1.32 | 1.18 | 2.50 | 17 |
| Dm | 0.07 | 0.19 | 0.13 | -0.62 | -0.53 | 0.32 | 0.20 | -0.30 | -0.10 | 6 |
| Eb | -0.18 | -0.35 | -0.14 | -0.75 | -0.63 | -0.03 | -0.32 | -0.78 | -1.11 | 2 |
| Lf | 0.35 | -0.06 | 0.38 | 0.51 | -0.57 | 0.65 | 0.73 | 1.15 | 1.89 | 11 |
| Mm | 0.51 | 0.34 | 0.50 | 0.56 | -0.07 | 0.55 | 1.01 | 1.10 | 2.11 | 15 |
| Pd | -0.03 | 0.52 | 0.51 | 0.27 | 0.16 | 0.02 | 0.48 | 0.28 | 0.77 | 9 |
| Pe | 0.26 | 0.51 | 0.06 | -0.05 | 0.33 | -0.33 | 0.32 | -0.38 | -0.06 | 7 |
| Po | 0.12 | 0.72 | 0.39 | 0.74 | 0.30 | -0.10 | 0.51 | 0.64 | 1.14 | 10 |
| Tb | 0.46 | 0.60 | 0.34 | 0.59 | -0.57 | 0.82 | 0.80 | 1.41 | 2.21 | 13 |
| Tf | -0.04 | -0.16 | 0.81 | 0.63 | -0.21 | 0.34 | 0.77 | 0.98 | 1.75 | 12 |
| Tg | -0.59 | -0.14 | 0.43 | -0.89 | -0.54 | 0.12 | -0.17 | -0.78 | -0.94 | 3 |
| Tm | 0.50 | -0.64 | 0.55 | 0.69 | 0.90 | -0.80 | 1.05 | -0.11 | 0.95 | 16 |
| Tv | 0.33 | 0.19 | 0.48 | 0.31 | -0.44 | 0.62 | 0.82 | 0.93 | 1.74 | 14 |
| Average | 0.22 | 0.15 | 0.21 | 0.34 | -0.12 | 0.20 | 0.44 | 0.54 | 0.98 |  |
| Range | 1.324 | 1.362 | 1.515 | 1.852 | 1.902 | 1.895 | 1.666 | 2.588 | 3.606 |  |
| B. CSI Table |  |  |  |  |  |  |  |  |  | Rain rank |
| Orchid | R-F CSI | R-P CSI | R-SV CSI | H-F CSI | H-P CSI | H-SV CSI | Rain CSI | Heat CSI | Total CSI |  |
| Ca | 0.74 | 0.96 | 0.11 | 0.37 | 0.45 | 0.11 | 0.85 | 0.48 | 1.33 | 3 |
| Cd | 0.46 | 0.76 | 0.27 | 0.56 | 0.16 | 0.30 | 0.73 | 0.86 | 1.59 | 2 |
| Cf | 1.18 | 0.71 | 0.20 | 0.34 | 0.45 | 0.30 | 1.38 | 0.64 | 2.02 | 11 |
| Cl | 1.00 | 0.91 | 0.34 | 0.15 | 0.91 | 0.97 | 1.34 | 1.12 | 2.46 | 9 |
| Db | 0.68 | 0.19 | 0.20 | 0.91 | 0.19 | 0.46 | 0.89 | 1.37 | 2.26 | 4 |
| Dm | 0.52 | 0.77 | 0.66 | 0.63 | 0.19 | 0.33 | 1.17 | 0.96 | 2.13 | 7 |
| Eb | 0.87 | 0.51 | 1.01 | 0.70 | 0.18 | 0.54 | 1.88 | 1.24 | 3.13 | 14 |
| Lf | 0.63 | 0.77 | 0.49 | 0.39 | 0.60 | 0.40 | 1.12 | 0.79 | 1.91 | 6 |
| Mm | 0.26 | 0.59 | 0.38 | 0.99 | 0.87 | 1.11 | 0.63 | 2.10 | 2.73 | 1 |
| Pd | 0.63 | 0.52 | 0.45 | 0.86 | 0.29 | 0.63 | 1.07 | 1.48 | 2.55 | 5 |
| Pe | 1.34 | 0.45 | 0.91 | 0.85 | 0.68 | 0.68 | 2.25 | 1.54 | 3.79 | 17 |
| Po | 0.65 | 0.18 | 0.71 | 0.20 | 0.69 | 0.69 | 1.36 | 0.88 | 2.24 | 10 |
| Tb | 1.10 | 0.32 | 0.83 | 0.10 | 0.86 | 0.65 | 1.94 | 0.75 | 2.69 | 15 |
| Tf | 0.71 | 0.91 | 0.51 | 0.24 | 0.37 | 0.37 | 1.23 | 0.61 | 1.84 | 8 |
| Tg | 0.55 | 0.47 | 0.93 | 0.55 | 0.27 | 0.38 | 1.48 | 0.93 | 2.41 | 12 |
| Tm | 1.08 | 0.56 | 0.89 | 0.24 | 0.38 | 0.51 | 1.97 | 0.75 | 2.71 | 16 |
| Tv | 0.46 | 0.27 | 1.23 | 0.68 | 0.30 | 0.56 | 1.69 | 1.24 | 2.93 | 13 |
| Average | 0.76 | 0.58 | 0.60 | 0.52 | 0.46 | 0.53 | 1.35 | 1.04 | 2.40 |  |
| Range | 1.08 | 0.78 | 1.12 | 0.89 | 0.75 | 1.00 | 1.62 | 1.62 | 2.45 |  |
| Units: | Flower no. | P / F | mm3 / spike | Flower no. | P / F | mm3 / spike | F+SV | F+SV | Rain+Heat |  |
| Labels: | R = rain, H = heat, F = flowering, P = pollination, SV = seed volume |  |  |  |  |  |  |  |  |  |

Comparisons of Index values in Table 3 show that flower production and seed yield (volume) were most informative overall and could be combined as rain or heat CRIs and a unified CRI for rain and heat (Table 3). Response (CRI), sensitivity (CSI) and combined (CI) indexes based on rain or heat ranked climate tolerance of species in a different order in Figure 13, but CRIs were most powerful overall (Table 3).

##### Box 1. Climate index analysis framework.

1. Collate local climate data for seasons and species productivity data for the same years (Table 2A).
2. Assess data by direct visualisation using graphs and statistical tests of significance, which requires adequate replication as determined by the number of years of data (e.g. Figs 7-10).
3. Produce correlation matrixes from data (e.g. seasonal rainfall vs. flower production), as shown in Table 2B.
4. Select maximum and minimum seasonal correlations for each species (Table 2C).
5. Use minimum and maximum correlation values to calculate climate response (CRI), sensitivity (CSI) and combined (CI) indexes (Table 2C).
6. Compile climate index tables by contrasting all pairs of productivity and climate data (Table 2D).
7. Compare the effectiveness of alternative climate or productivity data and index types for classifying climate tolerance of species (Table 3, Fig. 13).
8. Summarise climate effects relative to seasons (Fig. 12) and species (Section 5).
9. Assign climatic susceptibility ranks to species based on a single index (Table 3D), multi-index ordinations, or indexes contrasted with another functional trait (see Section 5 and 6).
10. Use regression or modelling approaches to forecast climate effects on species relative to past or possible future climate regimes (see Section 7).
11. Summarise data in vital statistics tables for species (Table S5, Brundrett 2016) including other key traits for reproduction, demography, fire responses, etc. (Table S6, Brundrett 2025).
12. Develop long-term monitoring frameworks and management plans for species and their habitats (Sections 9, 10).

Figure 14 compares the magnitude of CRIs for orchid species relative to temporal scales used to summarise climate data. Comparisons based on conventional seasons primarily show variations between species as positive effects from increased rain and negative effects from higher temperatures. As in Figure 11, there were similar results using conventional and two-month Noongar seasons (compare Fig. 14A, B to C, D), but CRIs from monthly climate data were less-well correlated with other results (Fig. 14E, F). Overall, 11 species had positive reposes to rain, three had negative responses and two had mixed reposes. Regarding temperature, 8 species had negative responses, 2 had weak positive responses and 7 had mixed responses to warmer conditions (Fig. 14B, D). Overall, differing responses to rainfall or heat were common, but contrasting flower and seed production responses occurred in fewer species, most of which had visually deceptive pollination.

**Figure 14.**
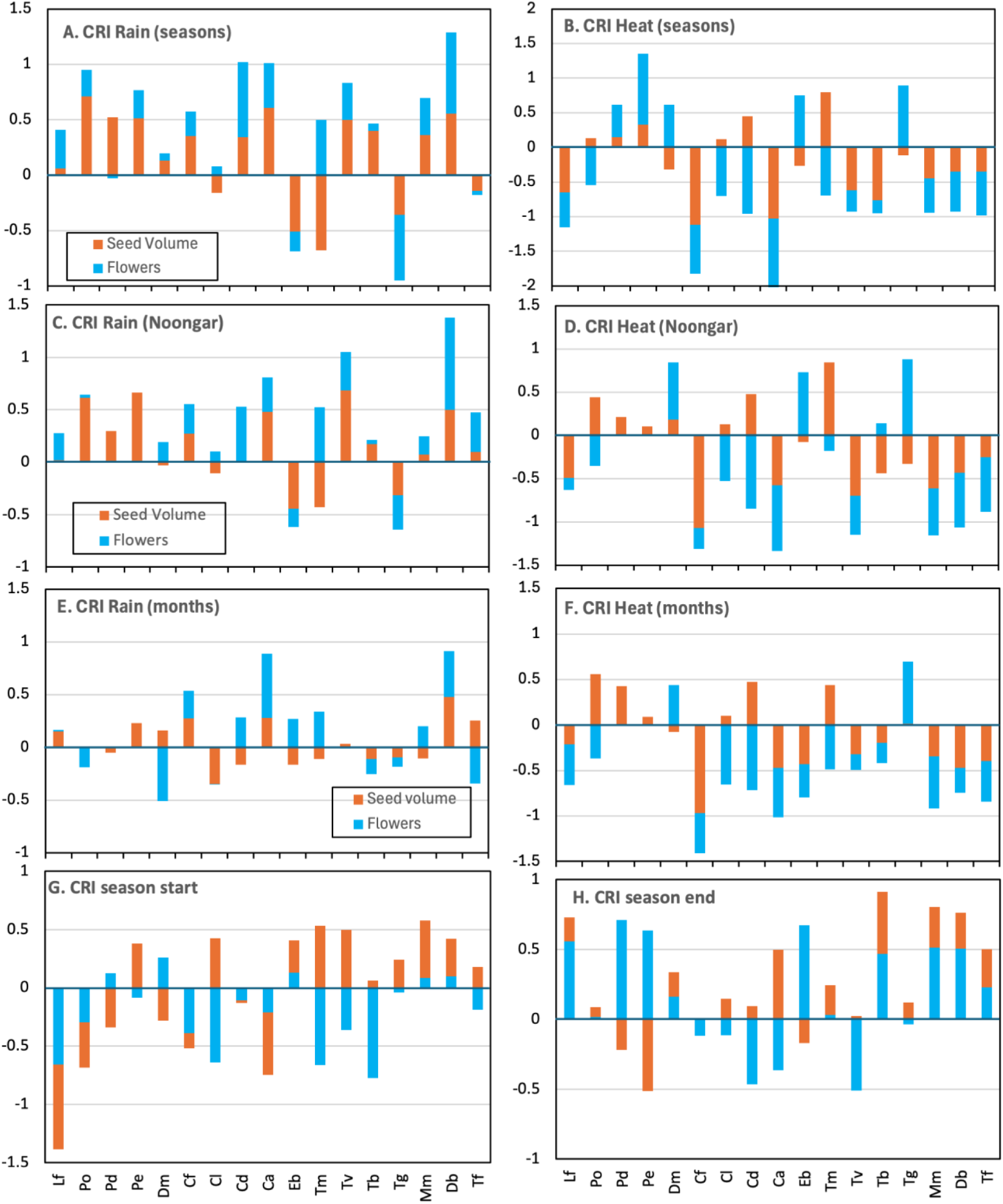
The magnitude and direction of the Climate Response Indexes (CRI) showing rain (**A, C, E**) or temperature (**B, D, F**) effects on 17 orchids arranged in flowering sequence (see Table 1 for species names). Climatic data are compiled using conventional seasons (3 monthly – **A, B**), approximated Noongar seasons (2 monthly – **C, D**), or monthly records (**E, F**). Correlations with growing season start (**G**) and end (**H**) dates are also presented.

Sorting CRIs by orchid phenology in Figure 14 revealed that earlier flowering orchids (autumn to early spring) were substantially more likely to experience negative effects from drought (Fig. 14A, C). Late flowering species were most likely to have negative responses to higher temperatures, but so did many that flowered earlier (Fig. 14B, D). Early flowering orchids were also impacted most by growing seasons delayed by autumn drought, while late-flowering species benefit from late spring rain that extended growing seasons (Fig. 14G, H).

### 5. Ranking species susceptibility

Climate-productivity correlations for orchids can be used to compare seasonal trends (Fig. 12), productivity measures (Fig. 14) or impacts on species (Fig. 15). Indexes could also be combined as ordination axes, as in Figure 15A, B, which used rain vs. heat CRIs or CSIs to show divergent results from flowering or yield data. These ordinations show a weaker trend for seed production than flowering for CRIs, but the reverse occurred for CSIs. An alternative approach used rain and heat indexes (with flower and seed production combined) as ordination axes for climate tolerances of species (Fig. 15C, D). This approach also sorted orchids by pollination syndromes, especially for CRIs (Fig. 15C), providing evidence of trait coordination (see Section 6).

**Figure 15.**
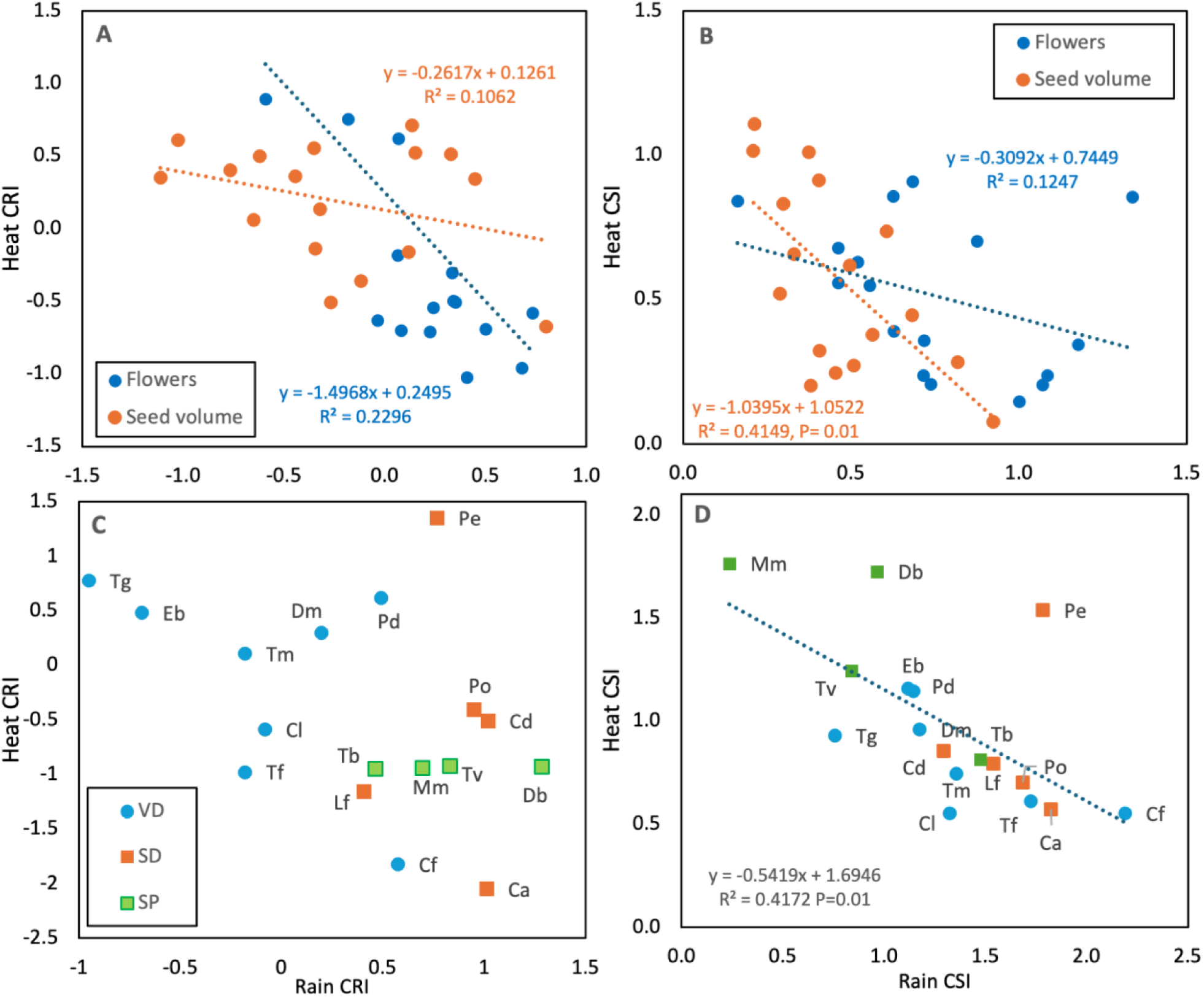
Categorising orchids using Climate Response (**A**) or Sensitivity (**B**) Indexes (CRI, CSI) for maximum temperature (heat) and rain for flower or seed production (Table 3). A separate approach contrasts rain or heat CRIs (**C**) or CSIs (**D**) using combined flower and seed production. Labels assign orchid pollination syndromes (VD = visual deception, SD = sexual deception, SP = self-pollination, see Table 1 for names, regressions and significant P values are shown).

As shown in Figure 16A, combined CRI values can be used to rank climate tolerance of species using data from Table 4. An alternative approach in Figure 16B combines direct responses (Figs. 7-10) with CRI analyses for rain and heat combined. Both approaches in Figure 16 rank species in a similar order, despite three species with strong inverse effects for rainfall and temperature (Fig. 16A) and five species with diverging scale and CRI ranks (Fig. 16B). Conflicting CRI results for species were also caused by differing climatic effects on flower or seed production (Fig 14).

**Figure 16.**
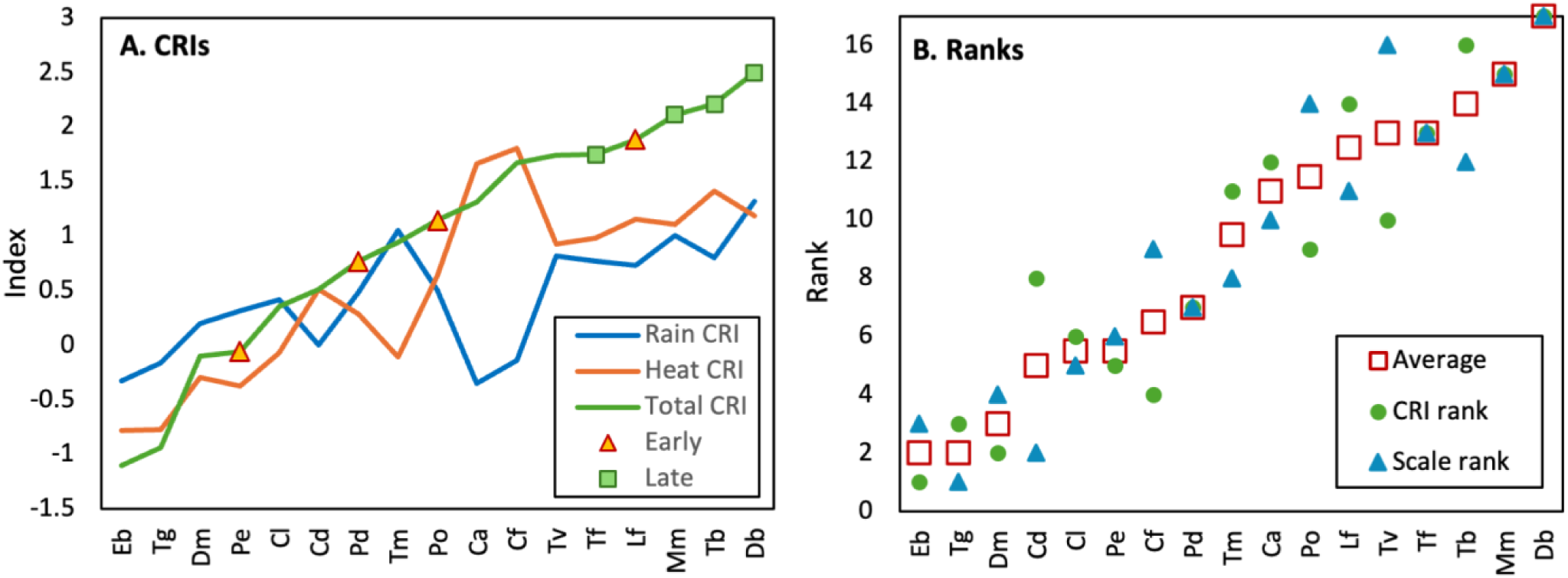
Ranking orchid climate responses using climate response (CRI) values for rain, heat or combined, with early or late flowering species indicated (**A**). Comparing climate indexes and a categorical scale for ranking orchid susceptibility (**B**), as explained in the text (species names in Table 1).

**Table 4.** Climate response summary with categories for correlations and CRIs (Fig. 11, Table S4).

|  | A. Categorical correlation summary |  |  |  |  |  |  | B Climate index summary |  |  |  |  |  |  |  |
| --- | --- | --- | --- | --- | --- | --- | --- | --- | --- | --- | --- | --- | --- | --- | --- |
| Name | Rain-flower | Rain-seed | Heat-flower | Heat-seed | Rain | Heat inverse | Total | Scale rank | Rain CRI | Heat CRI | Total CRI rank | Rain CRI rank | Scale & CRI rank | Climate preferences | Climate impacts |
| *Caladenia arenicola* | 1 | 2 | 0 | -5 | 3 | 5 | 8 | 16 | -0.35 | 1.66 | 10 | 1 | 10 | Wet autumn, dry winter, cool all seasons | Strong, very conflicting |
| *Caladenia discoidea* | 2 | 0 | 0 | 1 | 2 | -1 | 1 | 5 | 0.00 | 0.51 | 6 | 5 | 4 | Warm dry winter | Moderate |
| *Caladenia flava* | 1 | 2 | -1 | 2 | 3 | -1 | 2 | 8 | -0.14 | 1.81 | 11 | 4 | 7 | Wet autumn, dry winter | Moderate, conflicting |
| *Caladenia latifolia* | 1 | -1 | -3 | 2 | 0 | 1 | 1 | 6 | 0.42 | -0.07 | 5 | 8 | 5 | Autumn rain | Moderate, conflicting |
| *Disa bracteata* | 3 | 2 | -1 | -3 | 5 | 4 | 9 | 17 | 1.32 | 1.18 | 17 | 17 | 17 | Cool wet spring | Very strong |
| *Diuris magnifica* | 0 | 0 | 4 | 5 | 0 | -9 | -9 | 1 | 0.20 | -0.30 | 3 | 6 | 3 | Warm/dry autumn and winter, wet spring | Limited, conflicting |
| *Elythranthera brunonis* | 0 | -2 | 2 | -1 | -2 | -1 | -3 | 3 | -0.32 | -0.78 | 1 | 2 | 1 | Warm dry seasons | Limited |
| *Leporella fimbria* | 0 | 1 | -1 | -3 | 1 | 4 | 5 | 11 | 0.73 | 1.15 | 14 | 11 | 12 | Cool wet autumn winter | Strong |
| *Microtis media* | 2 | 3 | -1 | -1 | 5 | 2 | 7 | 15 | 1.01 | 1.10 | 15 | 15 | 16 | Wet all seasons, cool spring | Very strong |
| *Pheladenia deflormis* | -1 | 3 | 0 | 1 | 2 | -1 | 1 | 7 | 0.48 | 0.28 | 7 | 9 | 8 | Wet autumn, warm winter, cool spring | Moderate, conflicting |
| *Pterostylis ectypha* | 3 | 1 | 0 | 1 | 4 | -1 | 3 | 9 | 0.32 | -0.38 | 4 | 7 | 6 | Wet autumn, warm dry winter | Moderate |
| *Pterostylis orbiculata* | 2 | 3 | -2 | 1 | 5 | 1 | 6 | 14 | 0.51 | 0.64 | 9 | 10 | 11 | Cool wet autumn, dry winter | Strong, conflicting |
| *Thelymitra benthamiana* | 1 | 2 | 0 | -2 | 3 | 2 | 5 | 12 | 0.80 | 1.41 | 16 | 13 | 15 | Wet autumn and spring, dry winter | Very strong, conflicting |
| *Thelymitra fuscolutea* | 1 | 0 | -2 | -2 | 1 | 4 | 5 | 13 | 0.77 | 0.98 | 13 | 12 | 14 | Wet autumn and spring, dry winter | Very strong, conflicting |
| *Thelymitra graminea* | -3 | -1 | -3 | 1 | -4 | 2 | -2 | 4 | -0.17 | -0.78 | 2 | 3 | 2 | Dry and cool winter and spring, | Limited |
| *Thelymitra macrophylla* | -2 | -2 | -2 | 4 | -4 | -2 | -6 | 2 | 1.05 | -0.11 | 8 | 16 | 9 | Warm/dry winter and spring, | Strong, very conflicting |
| *Thelymitra vulgata* | 1 | 1 | -1 | 0 | 2 | 1 | 3 | 10 | 0.82 | 0.93 | 12 | 14 | 13 | Wet cool winter | Strong, conflicting |
|  | 12 | 14 | -11 | 1 | 26 | 10 | 36 | Table S4 | Table 3 |  |  |  | Fig. 16B | Figs. 7-10, Table S4 |  |

High index values in Figure 16 and Table 4 identify five early or late flowering orchids as the most sensitive to drought overall. The four latest flowering orchids (*T. benthamiana, T. fuscolutea, M. media* and *D. bracteata*) were ranked within the top five for climate impacts. The first three orchids to flower (*L. fimbria, P. deformis* and *P. orbiculata*) were also relatively sensitive to climate extremes, especially *L. fimbria* which flowers in autumn and was ranked forth overall.

### 6. Ecological interactions with climate responses

Figures 15C, which uses rain and heat CRI values, shows strongly divergent climate responses for orchids based on their pollination syndromes. Similar results are observed in Figure 17 which plots orchid flowering phenology against climate response indexes. Rain-based CRIs separated orchids by pollination syndromes into divergent groups with distinct slopes and intercepts (Fig 17A-C), but temperature CRIs produced less distinct groups (Fig. 17D). These correlations provide evidence for convergent evolution for orchids sharing pollination traits and divergence from orchids with different syndromes. Overall, pollination syndromes had stronger correlations with climate tolerance than phenology, but the latter was also important, since most visually deceptive orchids flowered after sexually deceptive species, when insect activity tends to peak

**Figure 17.**
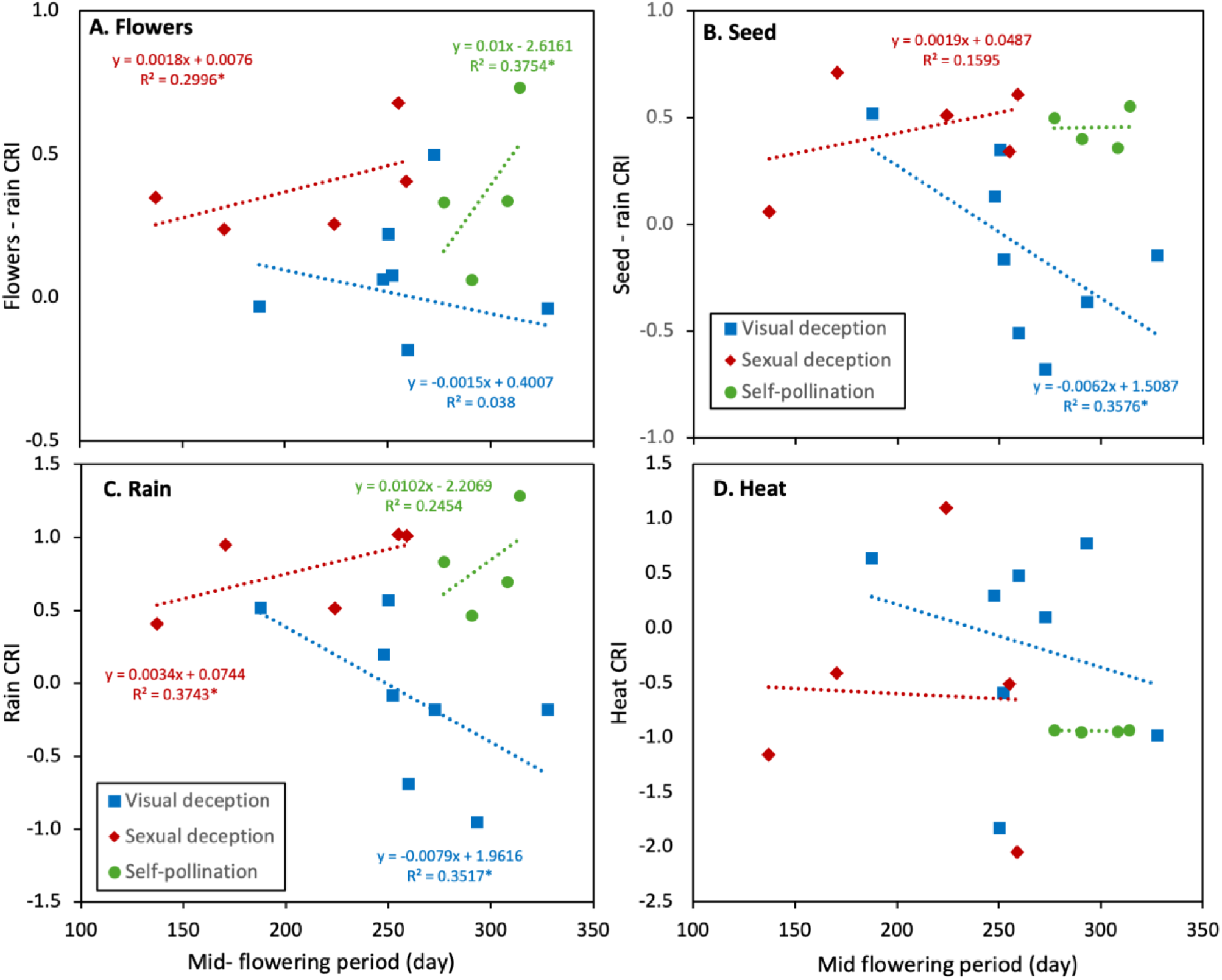
Comparing orchid flowering times and pollination syndromes with Climate Response Indexes (CRI) for seasonal rainfall effects on flower (**A**) or seed (**B**) production. Ordination using combined flower and seed CRIs are also shown for rain (**C**) or heat (**D**) (Significance values * = P < 0.1, ** = P <0.05, *** = P <0.001).

Four self-pollinating orchids formed a narrow group together near the middle of climatic ordinations (Fig. 15C) and in phenology-CRI analysis (Fig. 17). This response similarity is unexpected, since they belong to distantly related clades with closer insect pollinated relatives, suggesting that rapid convergent evolution of traits has occurred (O’Donnell et al. 2025). Another conundrum is the tendency for self-pollinating orchids to flower in late spring, which exposes them to hot dry conditions during seed production. This reduced yields, especially for *D. bracteata*, an invasive species responding to environmental cues from its origin in South Africa. However, this orchid seems to have broader climate tolerance in Australia than South Africa (Konowalik & Kolanowska 2018). In contrast *M. media,* which is more drought tolerant, is part of a widespread species complex, with a sister species (*M. unifolia)* that extends from South Australia to Asia (its name changes at the WA border).

Figure 18 extends climate response analysis by contrasting rainfall CRI values with other key functional traits. This placed climate sensitivity into a wider context defined by traits regulating survival, spread and fire responses. In addition to pollination syndromes and phenology, the strongest climate response correlations were with orchid lifespans, flower production, flower and leaf size, fire sensitivity and tuber depth (Fig. 18A-E). Orchid Lifespans were only weakly correlated with climate tolerance (Fig 18A). However, over longer time-scales climate impacts are likely to be worse for short-lived orchids, due to reduced reproduction in years with dry seasons. Later-flowering orchids that were very climate sensitive, such as *T. benthamiana, T. fuscolutea*, *D. bracteata* and *M. media,* tended to produce smaller but more numerous flowers (Fig. 18B, C). These species produce up to 15 flowers per spike for sun orchids and >40 flowers for self-pollinating species (Table 1). Trade-offs between flower size and number suggest there is an advantage for producing many smaller flowers when pollination rates are inherently low or species do not require insects. Plant size, as represented by leaf length and number (Fig. 18D), is also significantly correlated with climate responses, with larger leaves associated with lower resilience. There also are significant relationship between climate and fire tolerance or tuber depth (Fig. 18E, F), which are strongly correlated with each other (Brundrett 2025). Thus, some orchids are very sensitive to both fire and climate, especially late flowering sun orchids and self-pollinating species (*D. bracteata* and *M. media*).

**Figure 18.**
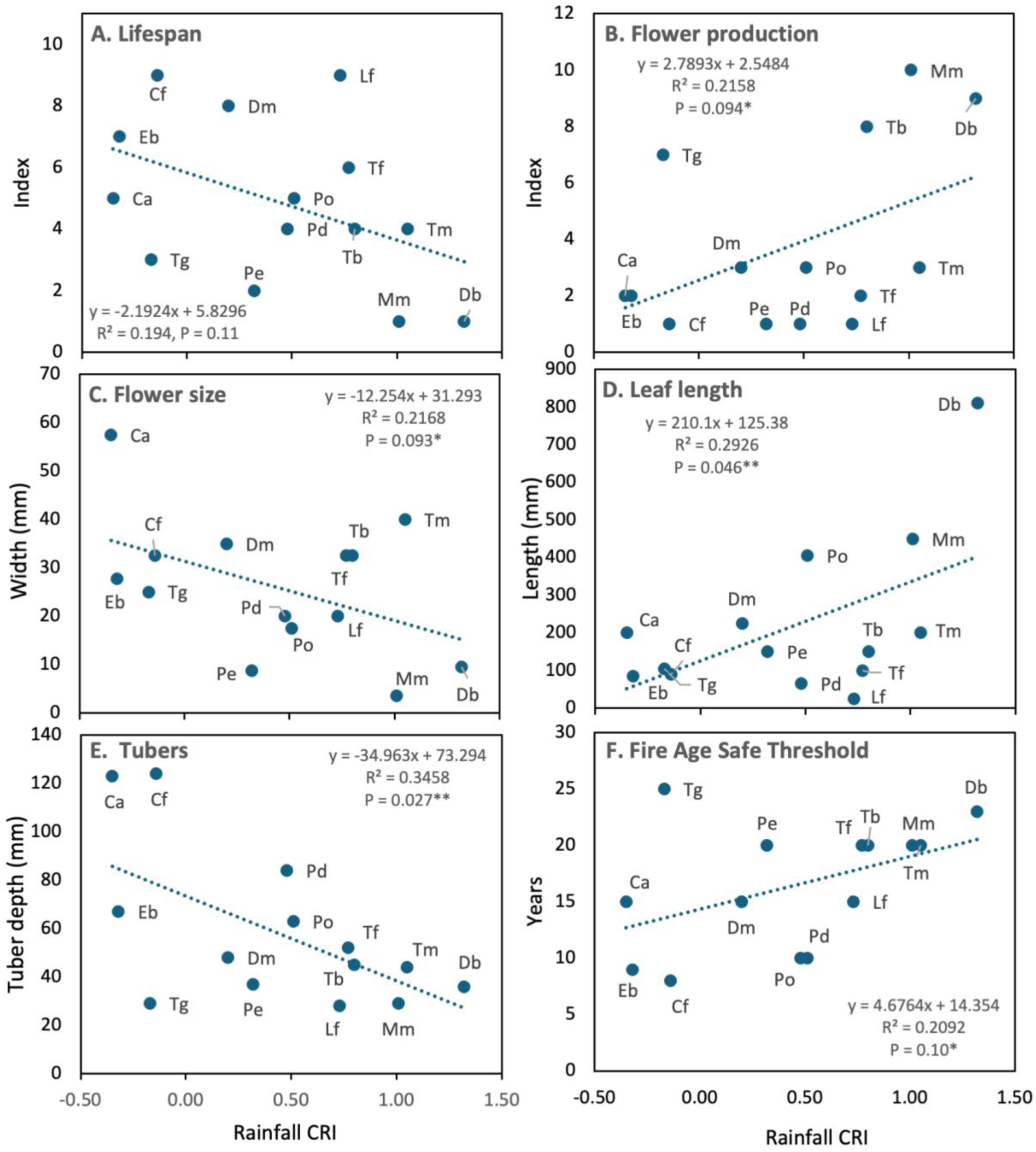
Correlations between orchid climatic or fire responses and other important traits (trait data from Table S6, species names in Table 1, * = P < 0.1, ** = P <0.05).

Taxonomic diversity of orchids compared here spans nine genera, including the three largest in Australia, and encompasses seven clades from three subfamilies (Diurideae, Cranichideae, Diseae). These orchids also have diverse phenological, reproductive and demographic traits, varying from highly clonal and long-lived to short-lived and seeding orchids, plus highly variable fire responses (Brundrett 2025). Correlations between climate responses and other important traits in Figure 18 reveal highly coordinated evolution between multiple axes in orchid trait space.

### 7. Climate impacts on insects, and fungi

Insects that pollinate orchids are also regulated by climate, with activity that varies seasonally and is very temperature dependant (Edens-Meier et al. 2013, Reiter et al. 2023, Kuiter 2025). *Thelymitra* flowers open on warm days (Edens-Meier et al. 2013) and bees that pollinate visually deceptive orchids also prefer warm weather (Kuiter 2025, Scaccabarozzi et al. 2020). Visually deceptive orchids include sun and donkey orchids (*Thelymitra* and *Diuris*) attract bees using complex visual and olfactory signals (Scaccabarozzi et al. 2023, 2025). Flowering of orchids must be synchronised with specific insects that emerge following the same climatic cues.

As explained above, climatic impacts on insects were strongly supported by distinct clustering of orchids by their pollination syndromes and flowering phenology (Figs. 15, 17). Orchids with visually deceptive pollination generally formed more seed during drier and warmer conditions (Fig 17). They also tended to flower later which increases the availability and diversity of pollinators by coinciding with peak flowering for bee pollinated flowers supplying pollen and nectar. Most *Thelymitra* species flower relatively late, especially *T. fuscolutea* (chestnut sun orchid) which forms seeds in early summer after its leaves have dried and capsules can mature on dead flower stalks (Brundrett 2019). In contrast, a winter flowering visually deceptive orchid (*P. deformis*) had very low rates of pollination except after fire (Brundrett 2025), presumably due to low temperature effects on pollinators. Orchids with sexually deceptive pollination remained productive in cool wet conditions (Fig. 15C, D). For example, *Pterostylis* species, which are pollinated by fungus gnats, usually flower in winter when fungi are fruiting (Fig. 6). Sexually deceptive pollination of *Caladenia* species by thynnine wasps was also relatively tolerant of cool-wet conditions (Fig. 9).

Some orchids respond to rain that falls before they break dormancy in winter (Figs. 8-11). These include *T. macrophylla* and *P. orbiculata* that form new subterranean shoots in autumn, as well as *C. arenicola* and *D. bracteata*, that emerge several months later. Autumn rain effects that occur before leaf growth are most likely due to recovery of soil physical or biological process after prolonged drought, and ultimately benefit orchids via their mycorrhizal fungi. Some orchids are also impacted by fungal pathogens associated damp winter conditions. These include frequently observed botrytis-like fungal disease that destroys leaves and flowers of *P. orbiculata* and rust fungus pustules in leaves of *T. fuscolutea* (Brundrett 2019). Rates of grazing, especially by invertebrates and rabbits, also varied considerably from year to year. Thus, there is a complex and delicate balance between conditions that favour orchids, their pollinators and their mycorrhizal fungi, as well as their parasites and grazers, all of which would be affected by climate extremes.

### 8. Extending climatic analyses across space, time and diversity

Orchids may respond to climate change by adjusting their periods of growth and flowering. For example, widespread species flower earlier in the north and later in the south of WA, following the progression of peak flowering of other native plants (Brundrett 2014). Herbarium records (www.ala.org.au) show that flowering phenology is relatively consistent, but autumn flowering species now appear later than in the past (e.g. *L. fimbria*). I also noted that flowers that open first or last on spikes were more likely to be pollinated (*L. fimbria* or *T. macrophylla*), but more research comparing chronologies of insects and orchids is required.

Climatic responses presented here are based on orchid performance in recent years, which may be substantially different from the past. Recent studies in WA found similar overall pollination rates to Table 1 for species of *Caladenia, Diuris* and *Pterostylis* (Elliott and Ladd 2002, Newman et al. 2013, Brundrett 2018, Scaccabarozzi et al. 2020). Data for *L. fimbria* from 1984 to 1987 by Peakall (1989) from sites nearby were also similar to the research site results 40 years later (Brundrett 2019). In contrast, Rica Erickson (1951) observed far higher insect visitation rates about 100 years before the current study. She noted that a *Diuris* species closely related to *D. magnifica* and *Caladenia flava* had pollen removed from every flower and most had pollen deposited by native bees. This historic trend is supported by Song et al. (2025), who found pollen removal and deposition declined after 1970 in herbarium specimens. Models based on future climate scenarios for *L. fimbria* by Kolanowska et al. (2021) found greater potential impacts on distributions of the pollinator than the orchid. However, this was based on the distribution of a single *Myrmecia* species (*M. urens*). This orchid is pollinated by several other species in eastern Australia (Kuiter 2025), and its range includes up to 50 *Myrmecia* species in WA (www.ala.org.au, 20-4-2026). *Leporella* pollinators probably include multiple species in WA that have not been identified.

Two case studies in Figure 19 extend climate response monitoring across the SWAFR bioregion. These show how geographic variation in rainfall provided very similar results to decadal comparisons at a single location for *D. bracteata* and *C. flava* (compare with Figs. 8 and 10). Flower and seed production by these two species was easily measured after flowering, especially for *C. flava* which I found in most botanical survey sites. *Disa bracteata* was also an ideal test subject, since it is a common weed that is drought sensitive and also reveals fire history (Brundrett 2025).

**Figure 19.**
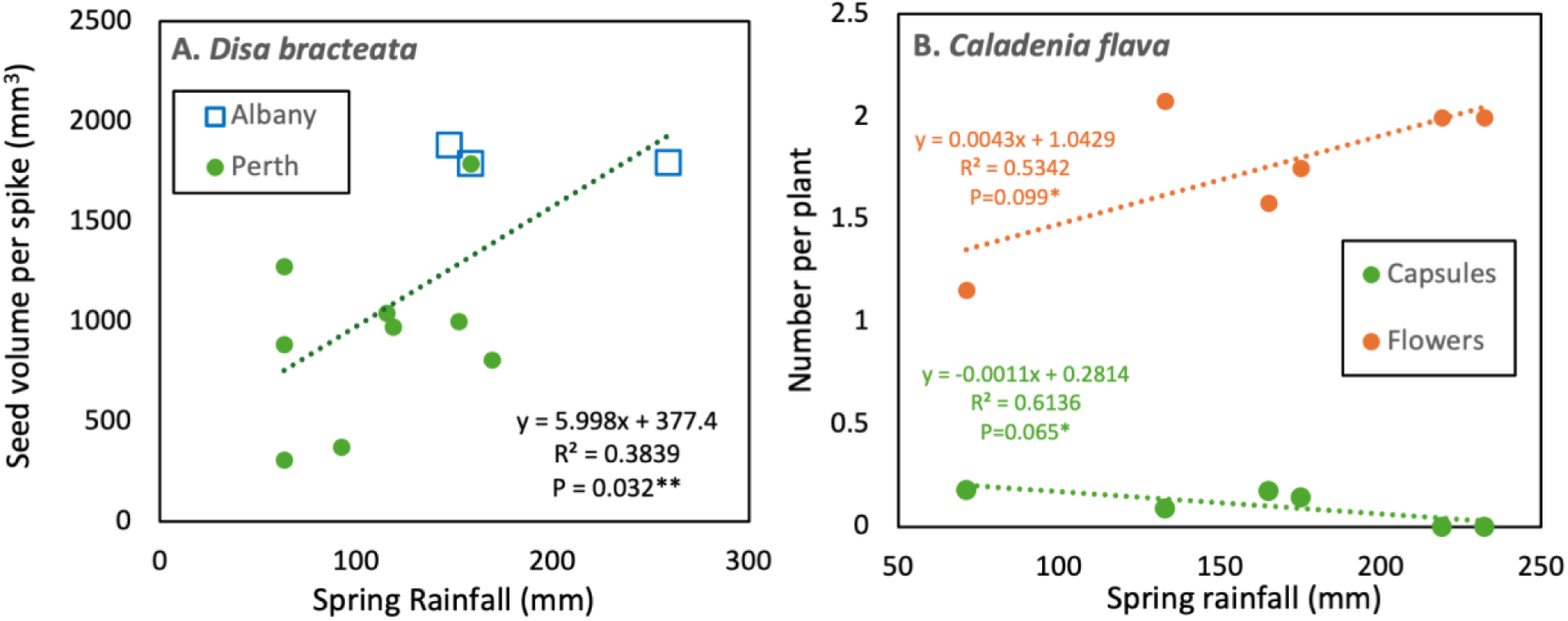
Case studies showing regional variations in rainfall effects on pollination (**A**) or flowering and pollination (**B**) for two widespread orchids. These were measured at two locations 450 km apart over several years (**A**) or six locations over a 600 km range in a single year (**B**).

Analyses based on CRIs can also be extended forward in time using extrapolated climate data. Figure 20 provides season climate forecasts based on past decadal rainfall or temperature trends (Fig. 1). These trends were used in combination with regression equations from productivity-climate analyses (Figs. 7-10) to provide climate impact predictions for eight species in Table 5A. Extrapolated effects of climate on flowering or seed production vary from 5% to 40% reductions, but one species (*T. macrophylla)* may have increased visually deceptive pollination (24%) due to drier and warmer conditions (Table 5). Case studies in Figure 20E-H uses data from Table 5 to illustrate how reductions in rainfall would be compounded by the increasing frequency of extremely dry years, especially in autumn. These graphs also compare three relatively sensitive species (Fig. 20E-G – *L. fimbria, P. deformis, D. bracteata*) to a less sensitive species (Fig. 20H – *C. arenicola*). Note that most species included in Table 5 are relatively responsive to climate so are suitable candidates for climate studies.

**Figure 20.**
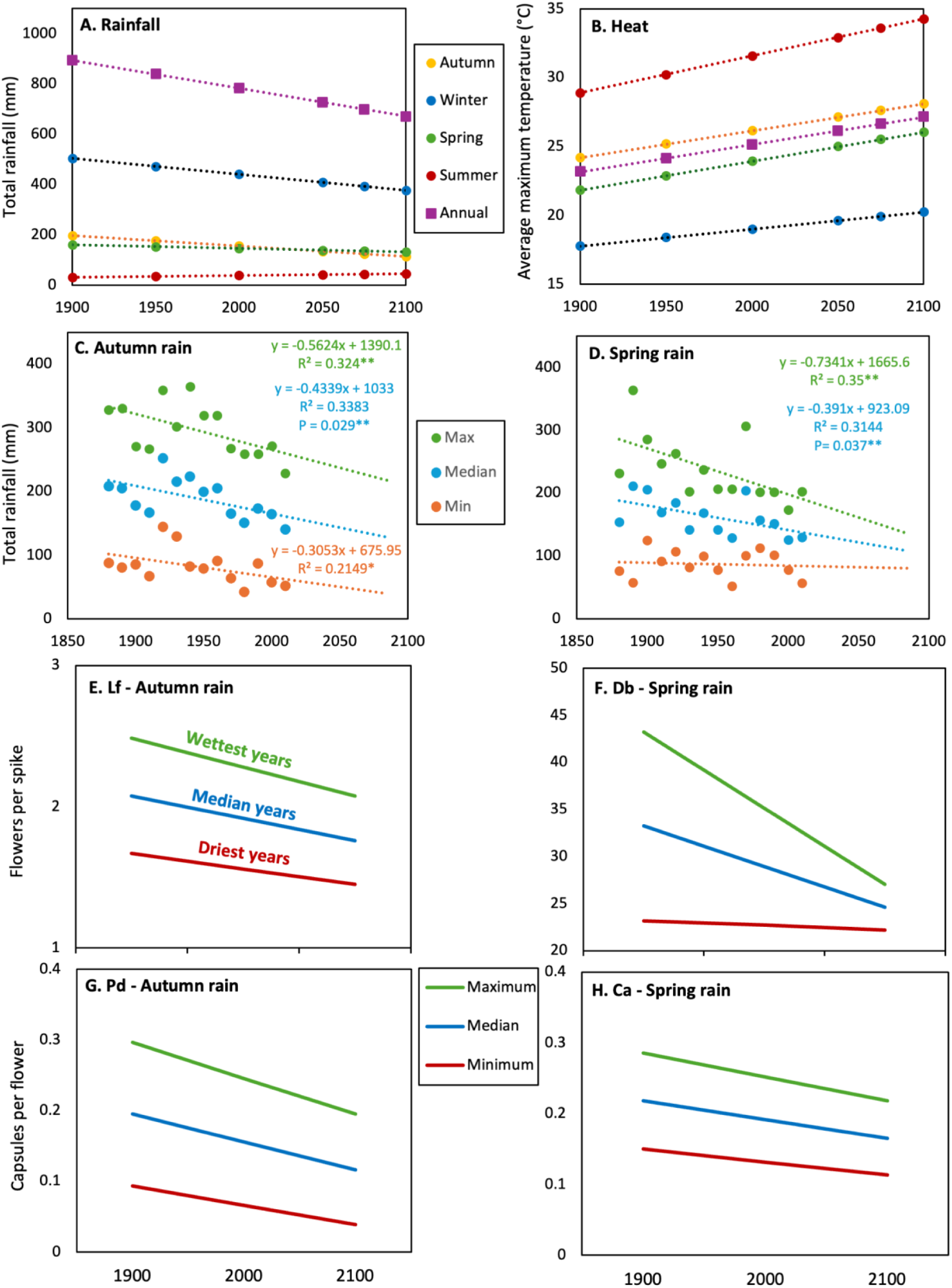
Seasonal rainfall deficit totals (**A**) and temperature averages (**B**) for Perth extrapolated to 2100 using data from Figure 1. Rainfall maximum, minimum and median collated by decade for autumn (**C**) or spring (**E**), extrapolated to 2100. Past and future impacts on orchid productivity are shown for extrapolated maximum, median and minimum predicted rainfall values for autumn (**E, G**) or spring (**F, H**). Results are for flower production (**E, F**) or pollination (**G, H**), and include highly climate sensitive (**E-G**) and a less sensitive orchid (**H**).

**Table 5.** Comparing recent and extrapolated future climate impacts on orchid productivity.

| A. Extrapolated impacts of seasonal rain on orchid flowering, pollination or seed production (1900-2100), |  |  |  |  |  |  |  |  | 200 years |  |  |
| --- | --- | --- | --- | --- | --- | --- | --- | --- | --- | --- | --- |
| Orchid | Factor | Response | R<sup>2</sup> | Intercept | Slope | 1900 | 2025 | 2100 | Change | % | Units |
|  | Autumn rain | Rainfall values |  |  |  | 197 | 146 | 115 | -82 | -42% | mm |
|  | Spring rain | Rainfall values |  |  |  | 153 | 143 | 133 | -20 | -13% | mm |
| Lf | Autumn rain | Flowering | 0.238 | 1.32 | 0.0036 | 2.03 | 1.85 | 1.74 | -0.30 | -15% | flowers/spike |
| Lf | Autumn rain | Pollination | 0.127 | 0.224 | 0.0008 | 0.382 | 0.341 | 0.316 | -0.07 | -17% | capsules/flower |
| Tm | Autumn rain | Flowering | 0.625 | 7.94 | 0.0602 | 19.8 | 16.7 | 14.9 | -4.94 | -25% | flowers/spike |
| Tm | Spring rain | Pollination | 0.223 | 0.0707 | -0.0003 | 0.025 | 0.028 | 0.031 | 0.006 | 24% | capsules/flower |
| Tm | Spring rain | Flowering | 0.205 | 9.83 | 0.0427 | 16.4 | 15.9 | 15.5 | -0.85 | -5% | flowers/spike |
| Db | Spring rain | Flowering | 0.501 | 13.3 | 0.1107 | 30.3 | 29.2 | 28.0 | -2.21 | -7% | flowers/spike |
| Tb | Spring rain | Seed yield | 0.282 | 504 | 1.084 | 670 | 659 | 648 | -21.7 | -3% | mm3 / spike |
| Ca | Autumn rain | Pollination | 0.471 | 0.0926 | 0.0006 | 0.211 | 0.180 | 0.162 | -0.05 | -23% | capsules/flower |
| Ca | Spring rain | Pollination | 0.337 | 0.0887 | 0.0006 | 0.207 | 0.175 | 0.169 | -0.04 | -19% | capsules/flower |
| Cd | Autumn rain | Flowering | 0.325 | 0.945 | 0.0051 | 1.95 | 1.69 | 1.53 | -0.42 | -21% | flowers/spike |
| Po | Autumn rain | Flowering | 0.428 | 3.34 | 0.0058 | 4.48 | 4.19 | 4.01 | -0.48 | -11% | flowers/spike |
| Pd | Autumn rain | Pollination | 0.272 | 0.0072 | 0.0009 | 0.185 | 0.139 | 0.111 | -0.07 | -40% | capsules/flower |
| Calculated from regression equations (Figs. 7-10) and projected rainfall declines (Fig. 20). |  |  |  |  |  |  |  |  | Max | 24% | rare positive prediction |
|  |  |  |  |  |  |  |  |  | Min | -40% |  |
| B. Actual variations in orchid productivity from 2016 to 2025. |  |  |  |  |  |  |  |  | Ave | -14% |  |
| Orchid | Factor | Response | Min | Max | Max-min | Variation | Summary |  |  |  |  |
| Lf | Orchid yield | Flowering | 1.34 | 2.69 | 1.35 | 50% | Max | 91% |  |  |  |
| Lf | for all | Pollination | 0.107 | 0.434 | 0.327 | 75% | Min | 19% |  |  |  |
| Tm |  | Flowering | 0.03 | 0.177 | 0.147 | 83% | Ave | 65% |  |  |  |
| Tm |  | Pollination | 0.0056 | 0.064 | 0.0584 | 91% |  |  |  |  |  |
| Tm |  | Flowering | 0.03 | 0.177 | 0.147 | 83% |  |  |  |  |  |
| Db |  | Flowering | 14.6 | 37.3 | 22.7 | 61% |  |  |  |  |  |
| Tb |  | Seed volume | 529 | 810 | 281 | 35% |  |  |  |  |  |
| Ca |  | Pollination | 0.0465 | 0.231 | 0.1845 | 80% |  |  |  |  |  |
| Cd |  | Flowering | 1.048 | 2.64 | 1.592 | 60% |  |  |  |  |  |
| Po |  | Flowering | 3.84 | 4.77 | 0.93 | 19% |  |  |  |  |  |
| Pd |  | Pollination | 0.027 | 0.136 | 0.109 | 80% |  |  |  |  |  |

Values for climate indexes provided here are likely to remain relevant in the future because they are based on a decade of high climatic variability where substantial impacts on orchids occurred, that may equal or exceed trends predicted in the future (Table 5B). Summarising climate history by decade revealed that the number of extremely dry autumns per decade increased faster than median rainfall trends, while rainfall totals for the wettest years decreased fastest in spring (Fig. 20C, D). Thus, productivity losses due to forecast average rainfall declines of 145 mm in autumn and 125 mm in spring would be compounded by the impacts of increasingly extreme climate anomalies (Table 5A, Fig. 20A, B). Contractions in growing season length are also very important, but require investigation. Research is urgently required to calculate critical climate thresholds for species and map spatial variations in predicted climate anomalies (Section 10). Extrapolations in Table 5 primarily concern orchids, but impacts on essential symbiotic insects and fungi are also very likely.

Analysis of climatic effects on species is likely to be complicated by overriding factors such as plant density, habitat condition and fire history (Collins & Brundrett 2015, Brundrett 2016, 2025). Climate anomalies and fires also have major impacts on orchid habits (e.g. canopy cover, litter, weeds). Despite all these substantial impacts, most orchids in this study have relatively stable or expanding local populations, but there are concerning trends for some species, especially if they are also fire intolerant (Fig. 18E).

The applicability of climate indexes to plants other than terrestrial orchids and organisms in other kingdoms requires investigation, as does the use of additional productivity measures and climatic variables. Note that the inclusion of more categories of climate data may have limited benefits, since CRI analysis only requires the highest and lowest correlations with productivity.

### 9. Orchid climate, fire and habitat observatory networks

This study revealed substantial impacts of climate change on orchids from a localised decadal study and short-term geographic comparisons. Both approaches can be used establish “Orchid Climate Observatories”, as defined in Figure 21. These living laboratories monitor effects of local climate conditions on orchid populations, their pollinators and habitats (e.g. canopy cover, weeds, grazing). Some orchids make very good indicators of fire impacts, by measuring their abundance and diversity relative to fire age or contrasting recently burnt to long unburnt areas (Brundrett 2025). Orchid observatories would function as educational activities, research networks and citizen science projects, which aim to provide essential data for orchid ecology, sustainability and conservation.

**Figure 21.**
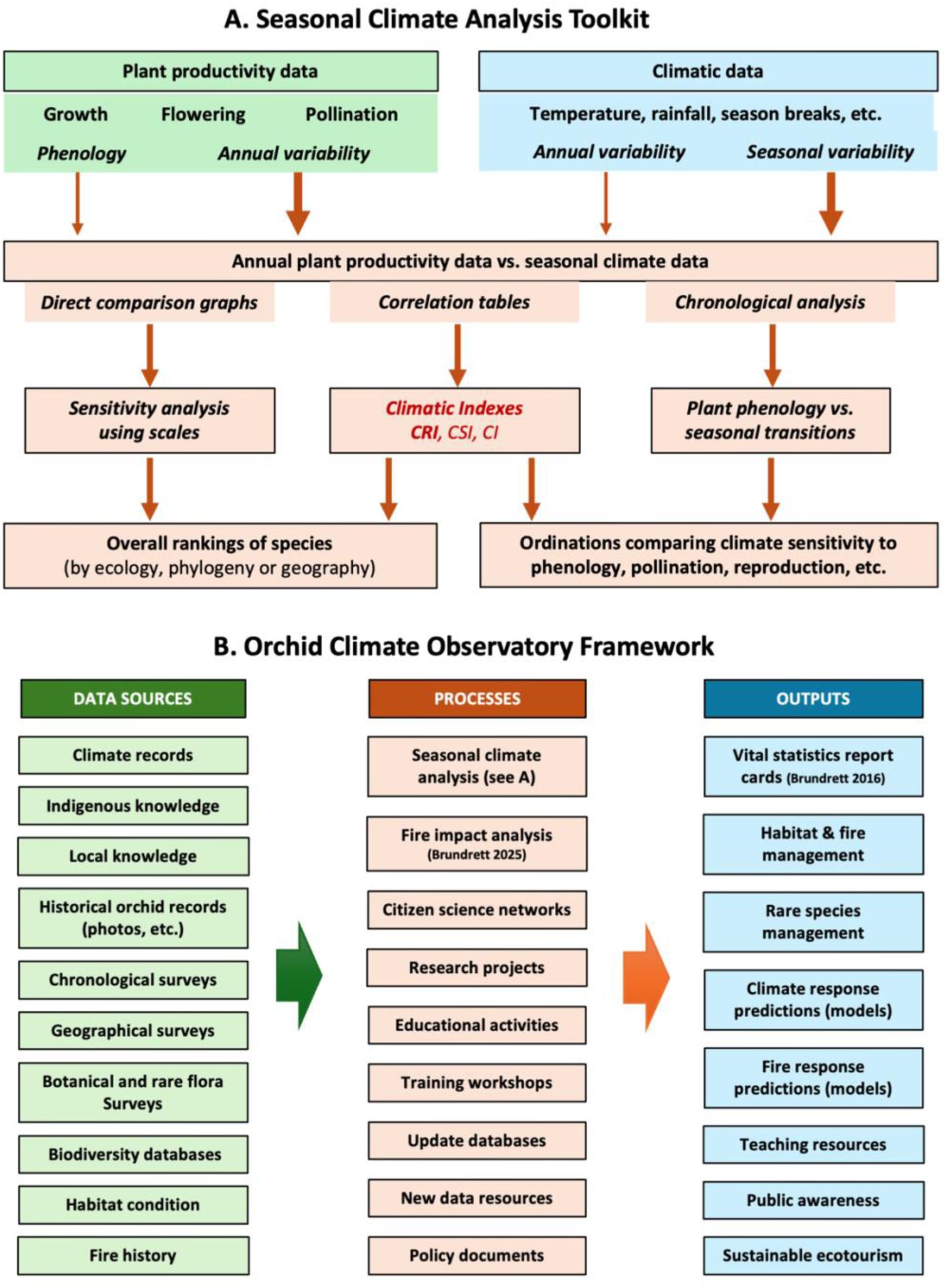
Workflows for climate analysis (A) and climate observatories (B) using orchids as an example.

Orchids can be very climate sensitive so functioning like “canaries in a coal mine”. These iconic species, have enthusiast networks that observe the same areas annually, and take photographs documenting flowering times, pollination and habitat condition spanning decades. They could also record orchid flowering and seed production, which is relatively quick and easy relative to costs and time expended to visit locations. Volunteers can also record threats to orchids and provide local climate records resolving spatial variability and missing data (Fig. 21B). Other potential observations include the abundance of other flowers that feed orchid pollinators, which may have differing climatic susceptibilities (Kolanowska & Scaccabarozzi 2024), and the relative abundance of native bees and honeybees - which can cause severe competition and be ineffective pollinators (Scaccabarozzi et al. 2024). Direct observations of orchid pollination is also extremely valuable but more time consuming (see Kuiter 2025). If possible, monitored orchids should include high and low climate resilience, be relatively common and widespread, are readily identifiable (this excludes many WA orchids), and represent different pollination syndromes and phenologies (see examples in Table 5). It would be expected that relatively widespread and common species are more adaptable to climate change than uncommon species, but this requires confirmation. It might also be expected that widespread orchids are less likely to adapt to local conditions due to their capacity for long range wind dispersal, but that also requires investigation. This is less relevant when studying responses to rapid changes in climate, when local and widespread sampling have similar results (Fig. 19).

### 10. Species and ecosystem management

Climate and fire monitoring tools presented here (Fig. 21) are applicable to the management of rare species and their habitats, especially when risks from altered climate of fire regimes are suspected. This approach can be extended to include other key traits. These can be summarised in vital statistics report cards (Brundrett 2016), as demonstrated for three very rare SWAFR orchids in Table S5. A trait assessment toolkit is presented in Table S6, which provides definitions and methodologies for measurement of key ecological properties and processes, including fire response and climate metrics (Brundrett 2025). Knowledge of orchid climatic and fire sensitivity is most critical for rare species, but is often lacking (Brundrett 2016). Orchid climate observatories could also be a major resource for determining orchid population viability and changes to habitat conditions (Fig. 21B).

## Conclusions

This is the third comprehensive study located in the same urban orchid diversity hotspot (Brundrett 2019, 2025), confirming the value of research on common species in accessible locations. Rainfall reductions and temperature increases became more severe and frequent over 125 years at this site and were extremely variable during this study. This seasonal climate data was combined with orchid productivity data in a new climatic analysis workflow that measured responses of species. As explained in Figure 21, CRI analysis is a comprehensive toolkit based on flexible, robust and uncomplicated methods, and data that was easily collected non-destructively. This approach efficiently ranked climate sensitivities and tolerances of orchid species (Fig. 16). Rainfall deficits caused the greatest impacts on orchids in autumn and spring, resulting in shorter growing seasons, but visually deceptive pollination could benefit from warm dry conditions. Potential impacts on pollinators and mycorrhizal fungi were also detected and reductions in habitat quality, including canopy loss and fire impacts were observed. Thus, knowledge of both direct and indirect climate impacts on orchids is required for conservation and habitat management, and can be gained from local climate laboratories (Section 10).

Climate indexes sorted taxonomically and functionally diverse orchids into susceptibility gradients that were strongly correlated with key traits like pollination, phenology, plant size, lifespan and fire tolerance. Extreme functional trait complexity is widespread in the SWAFR flora, especially for pollination, fire and nutrition traits (Lambers et al. 2010, Brundrett 2021, Brundrett et al. 2024, 2025). However, SWAFR orchids provide some of the most extreme examples of evolutionary trends leading to trait complexity, integration and divergence or convergence. It has yet to be determined if this complexity extends to orchids found elsewhere. Other knowledge gaps include: (i) detailed modelling of future climate responses, (ii) measuring separate impacts on pollinators and mycorrhizal fungi relative to orchid responses, (iii) applying CRI analysis to orchid conservation and (iv) extending it to other geographic regions and wider biodiversity. Orchids are effective model species in climate observatories, which are based on straightforward and adaptable workflows (Fig. 21). This framework should also be relevant to other plants, including those grown for agriculture and horticulture, and members of other kingdoms, by utilising existing climatic records along with productivity data that is readily collected or may already exist.

## Data Availability

The data that support this study are available in the article and accompanying online supplementary material.

## Conflicts of interest

The author declares no conflicts of interest.

## Declaration of funding

This research did not receive any specific funding.

## Acknowledgements

Special thanks to Karen Clarke for many walks through the study site and helping to discover local biodiversity. I am also very grateful to orchid enthusiasts who located orchids, The Friends of Warwick Bushland and City of Joondalup staff.

## Supplemental Tables

**Table S1.** Orchid productivity data for study period.

**Table S2.** Climate data for study decade.

**Table S3.** All climate-orchid productivity correlations and indexes

**Table S4.** Categorical summary of climate responses with CRI analysis summary

**Table S5.** Vital statistics report card categories for orchids (Brundrett 2016), extended to include fire and climate responses.

**Table S6.** Data sources and definitions for key orchid functional traits (Brundrett 2025), expanded to include climate analysis.

## Notes

### Competing Interest Statement

The authors have declared no competing interest.

### Summary of Updates

Minor corrections made to text for clarity and conciseness Hyperlinks fixed Figures standardised Tables are higher resolution

